# Structural basis of Cfr-mediated antimicrobial resistance and mechanisms for its evasion

**DOI:** 10.1101/2023.09.27.559749

**Authors:** Elena V. Aleksandrova, Kelvin J. Y. Wu, Ben I. C. Tresco, Egor A. Syroegin, Erin E. Killeavy, Samson M. Balasanyants, Maxim S. Svetlov, Steven T. Gregory, Gemma C. Atkinson, Andrew G. Myers, Yury S. Polikanov

## Abstract

The ribosome is an essential drug target as many classes of clinically important antibiotics bind and inhibit its functional centers. The catalytic peptidyl transferase center (PTC) is targeted by the broadest array of inhibitors belonging to several chemical classes. One of the most abundant and clinically prevalent mechanisms of resistance to PTC-acting drugs is C8-methylation of the universally conserved adenine residue 2503 (A2503) of the 23S rRNA by the methyltransferase Cfr. Despite its clinical significance, a sufficient understanding of the molecular mechanisms underlying Cfr-mediated resistance is currently lacking. In this work, we developed a method to express a functionally-active Cfr-methyltransferase in the thermophilic bacterium *Thermus thermophilus* and report a set of high-resolution structures of the Cfr-modified 70S ribosome containing aminoacyl- and peptidyl-tRNAs. Our structures reveal that an allosteric rearrangement of nucleotide A2062 upon Cfr-methylation of A2503 is likely responsible for the inability of some PTC inhibitors to bind to the ribosome, providing additional insights into the Cfr resistance mechanism. Lastly, by determining the structures of the Cfr-methylated ribosome in complex with the antibiotics iboxamycin and tylosin, we provide the structural bases behind two distinct mechanisms of evading Cfr-mediated resistance.

## INTRODUCTION

A preponderance of naturally evolved antibacterial agents and many synthetic drugs employed clinically for the treatment of human infectious diseases target the bacterial ribosome and thereby inhibit bacterial protein synthesis^1–3^. Among these, entire classes of chemically distinct antibiotics bind in or near the peptidyl transferase center (PTC), located within the large subunit of the bacterial ribosome, and often inhibit protein synthesis by direct competition with the binding of aminoacyl-tRNA (aa-tRNA) substrates. As a natural evolutionary response to antibiotic therapy, numerous antibiotic resistance genes have emerged in pathogenic bacteria, but among the more consequential and widespread of these genes is *cfr* (<u>c</u>hloramphenicol-florfenicol resistance), whose gene product – Cfr – is a methyltransferase that modifies the universally conserved ribosomal adenosyl residue A2503 of the 23S rRNA by C-methylation at position 8 (**Figure 1A**)^4,5^. This modification lies within the heart of the ribosome, near the A site of the PTC, and confers resistance to a wide range of PTC-targeting antibiotics, including phenicols, lincosamides, oxazolidinones, pleuromutilins, and streptogramins A (collectively known as PhLOPS_A_), as well as hygromycin A and 16-membered macrolides, which bind in the nascent peptide exit tunnel (NPET)^5–7^. Interestingly, the A2503 residue is also C-methylated at position 2 by the housekeeping rRNA-methyltransferase RlmN (**Figure 1A**), an enzyme widely distributed among bacteria^8^. RlmN and Cfr are homologs, sharing 34% sequence identity (*Escherichia coli* RlmN *versus Staphylococcus aureus* Cfr), and both employ S-adenosyl methionine as methyl donor^9^. RlmN-mediated C2-methylation of A2503 is constitutively present in 23S rRNA and contributes to translational fidelity^10–12^, whereas Cfr-mediated C8-methylation of the same nucleotide leads to multidrug resistance^5,9^.

**Figure 1.**
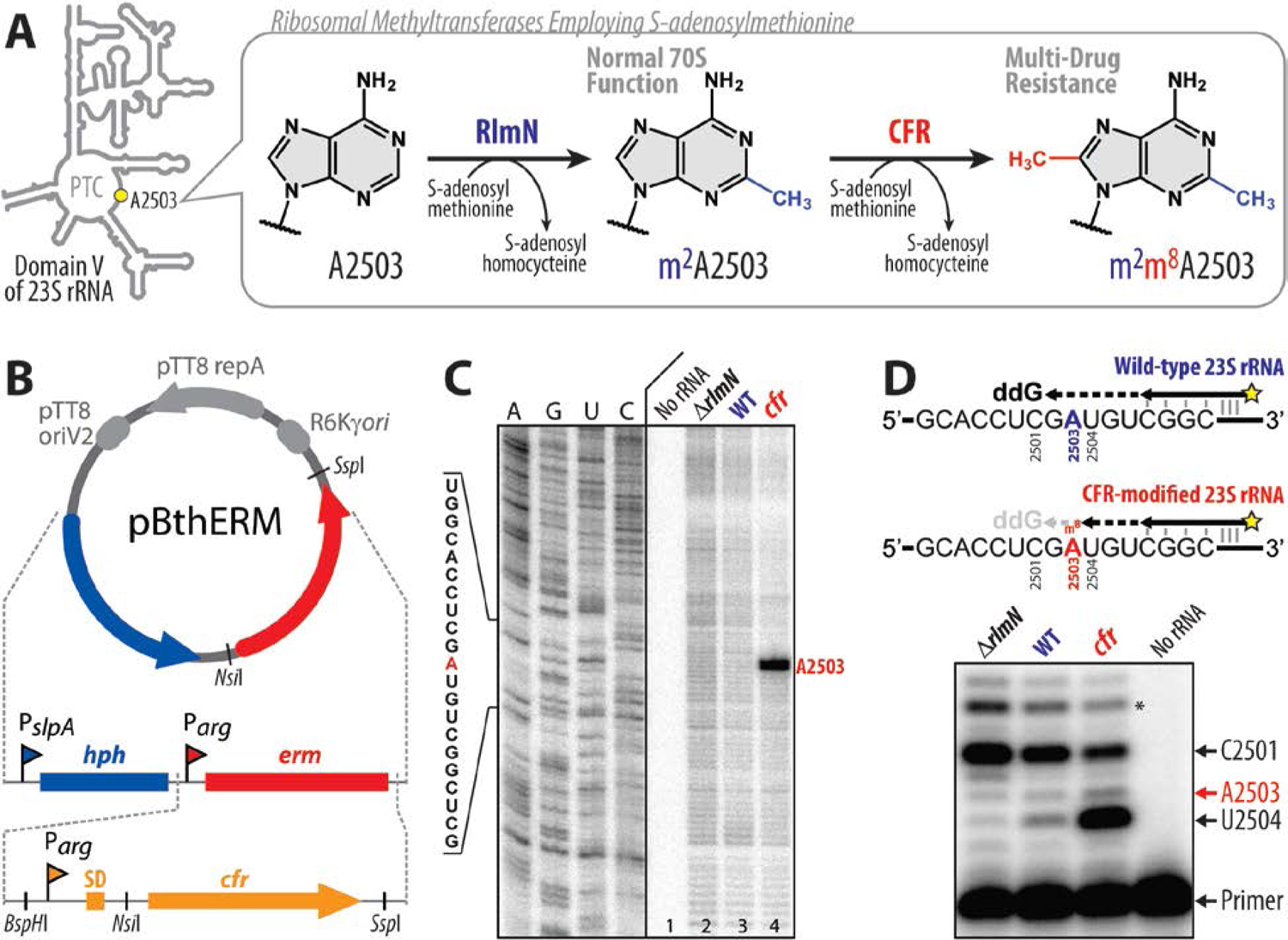
*Thermus thermophilus* HB27 strain expressing Cfr-like methyltransferase. (**A**) While C2-methylation of A2503 in domain V of the 23S rRNA by the housekeeping RlmN methyltransferase is required for normal function of the 70S ribosome, C8-methylation of the same residue by the Cfr-class methyltransferases results in high-level resistance to several chemically-unrelated classes of PTC-targeting antibiotics. (**B**) Schematic maps of the previous pBGAA1-BthERM vector (top) and its new derivatives with the *erm* gene (red) replaced by one of the six selected *cfr*-like genes (bottom) from moderately thermophilic bacteria. These vectors are designed to replicate in *Thermus thermophilus* (*Tth*). They carry the hygromycin B resistance marker *hph* under the control of P_slpA_ promoter (blue) and a *cfr*-like gene under the control of P_arg_ promoter (orange). The obtained vectors were used to express Cfr methyltransferases in *Tth* HB27 cells. (**C**) Primer extension analysis of 23S rRNA isolated from (Cfr+) *Tth* cells transformed with pBGAA1-PfuCFR vector (*cfr*) and control (Cfr-) wild-type *Tth* cells (WT). We used 23S rRNA isolated from the Δ*rlmN Tth* HB27 strain lacking A2503-C2-methylation (*ΔrlmN*) as an additional negative control. Reverse transcriptase stalls at C2,C8-dimethylated A2503 and produces truncated cDNA (marked with the red arrow). Sequencing lanes are shown on the left. (**D**) Precision primer extension analysis of the same 23S rRNA samples in the presence of dATP, dCTP, dTTP, and ddGTP. While the dideoxynucleotide ddGTP causes reverse transcription to stop at position C2501 on all templates, the Cfr-catalyzed C8-methylation of m^2^A2503 causes arrest of cDNA synthesis at the preceding position U2504 of the 23S rRNA. The extent of Cfr-methylation (∼70%) is calculated as the ratio of the intensity of the U2504-specific band to the sum of the intensities of the U2504-, C2501-specific and the readthrough bands (marked with an asterisk) after background subtraction. Experiments were repeated three times independently with similar results.

The plasmid-borne *cfr* gene was first discovered in 2000 in the bacterial species *Mammaliicoccus sciuri* (previously *Staphylococcus sciuri*)^13^ and was later found on plasmids in isolates from other sources^14^. In 2007, the first case of *cfr* occurrence in humans was identified in a clinical strain of methicillin-resistant *S. aureus* (MRSA, strain CM05), where the *cfr* gene was located on the chromosome^15^. Today, the *cfr* gene, with only minor sequence variations, has been found worldwide in pathogenic Gram-positive and Gram-negative bacteria isolated from humans and animals^16^.

Based on the recent structure of the drug-free Cfr-modified 50S ribosomal subunit from *E. coli*^17^, the observed effect of A2503-C8-methylation on the binding of some of the PTC-targeting antibiotics could be explained with a “*simple steric clash*” model. Alignments of this Cfr-modified ribosome with those of various ribosome-bound PTC inhibitors show that the newly introduced C8-methyl group in A2503 sterically overlaps with the binding sites of many classes of PTC-acting drugs. Therefore, it has been proposed that the Cfr-dependent A2503-C8-methylation is likely to interfere physically with antibiotic binding^17^. For some of these PTC-targeting antibiotics, the overlap with the C8-methyl group is small, raising the possibility of avoiding the clash via minor structural rearrangements of either the drug, the target site, or both. In principle, a collision between a drug molecule and the C8-methyl group can completely abolish binding, especially if the drug establishes only a few primary contacts with the ribosome, as is the case for chloramphenicol or linezolid. However, it is hard to rationalize this *simple steric clash* model for the 16-membered macrolides (such as josamycin or spiramycin), which form multiple strong interactions with the ribosome away from the C8-atom of A2503 in the PTC, yet exhibit decreased activity in *cfr*-expressing strains^6^. Thus, the prevailing *simple steric clash* model fails to adequately explain the observed Cfr-mediated resistance to some PTC-targeting antibiotics.

Alternatively, the mechanism of Cfr-mediated resistance, at least for some classes of drugs, could involve allosteric structural rearrangement(s) in the PTC that prevent drug binding only when aminoacyl- and/or peptidyl-tRNA ligands are present on the ribosome. The rationale for this hypothesis stems from the observation that key functional nucleotides around the PTC (A2062, U2506, U2585, A2602) change their positions upon binding of the tRNA substrates^18,19^. Importantly, one of these nucleotides (A2062) rotates relative to its position in a vacant ribosome in the presence of formyl-methionyl- or peptidyl-tRNA in the P site and forms a symmetric trans A-A Hoogsteen base pair with the residue A2503^20,21^. Thus, it is conceivable that Cfr-mediated C8-methylation of A2503 could allosterically affect the positions of other 23S rRNA nucleotides (such as A2062) in the presence of tRNAs, resulting in their conformations being incompatible with a drug binding to its functional site in the PTC. Although an attractive hypothesis, such Cfr-dependent structural rearrangements have not been observed to date.

To understand the structural basis of Cfr-mediated resistance, we first identified conditions for expressing a functionally-active Cfr-methyltransferase in the thermophilic bacterium *Thermus thermophilus* and solved the high-resolution X-ray crystal structure of the Cfr-modified 70S ribosome with non-hydrolyzable aa-tRNAs in both the A and P sites of the PTC. Using our recently developed approach for the semisynthesis of non-hydrolyzable peptidyl-tRNAs, we also solved the crystal structure of the Cfr-modified 70S ribosome with a peptidyl-tRNA in the P site. The new structures reported herein reveal an unexpected Cfr-induced displacement of nucleotide A2062, which is likely to be responsible for the inability of at least some PTC inhibitors to bind to the ribosome. Lastly, we determined the structure of the Cfr-modified 70S ribosome in complex with two antibiotics with activity against Cfr-expressing bacteria, iboxamycin and tylosin, and uncovered the structural bases behind their abilities to engage the Cfr-modified ribosome. Notably, we observe a displacement of the Cfr-methylated A2503 nucleotide by ∼1–1.5 Å upon binding of iboxamycin, which could not have been predicted on the basis of existing antibiotic-ribosome co-crystal structures.

## RESULTS

### Engineering of Cfr-expressing T. thermophilus strain

Cfr enzymes are expressed in a broad spectrum of pathogenic bacterial clinical isolates as well as in non-pathogenic bacteria, where they provide strong resistance to various PTC inhibitors^5,6,14–16^. As nucleotide A2503 of the 23S rRNA is inaccessible to enzymes in the mature 50S subunit, RlmN and Cfr methyltransferases must operate during the ribosome assembly proces^22,23^. Therefore, to isolate predominantly Cfr-modified ribosomes for crystallographic study, we first constructed a bacterial strain expressing a catalytically-active Cfr methyltransferase.

Based on our past work in obtaining Erm-modified 70S ribosomes (N6-dimethylated at position A2058 of the 23S rRNA) and solving its high-resolution structure^20^, we have chosen the same Gram-negative thermophilic bacterium *Thermus thermophilus* (*Tth*) as our experimental model. Because expression of a functionally active Cfr enzyme from the mesophilic bacterium *S. aureus* in the thermophilic bacterium *Tth* is unlikely to be successful, we selected five *cfr*-like genes from genomes of bacteria adapted for growth at elevated temperatures and used *S. aureus* Cfr as a control (**Figure S1, S2; Table S1**). The desired *cfr*-like genes were commercially synthesized, cloned into the previously assembled pBGAA1-BthERM vector (**Figure 1B**, top) to replace the *erm* gene^20^ (**Figure 1B**, bottom), and expressed in *Tth* HB27 cells.

Cfr expression in the resulting strains was assessed by a microbiological approach that exploits the sensitivity of WT *Tth* cells to several PTC-targeting drugs, such as chloramphenicol (CHL), florfenicol (FFL), lincomycin (LNC), clindamycin (CLI), or iboxamycin (IBX) (**Table S2**, blue). Antibiotic susceptibility testing of the plasmid-transformed *Tth* cells showed that the expression of a *cfr*-like gene from *Planifilum fimeticola* (referred hereafter as PfiCFR) or *Planifilum fulgidum* (PfuCFR), both Bacillota (firmicute) bacteria in the Thermoactinomycetaceae family, resulted in a substantial increase of MICs for CHL, FFL, LNC, and CLI by 32, 64, 1024, and 4096 fold, respectively (**Table S2**, red) but not for the NPET-targeting macrolide erythromycin (ERY), when compared with control cells carrying the empty vector (**Table S2**, green). This specific resistance pattern is unlikely to arise from spontaneous mutations (**Table S2**).

To verify that the observed drug resistance arises from the specific methyltransferase activity of PfuCFR, we used a primer extension analysis to assess A2503 modifications biochemically. This method is based on the arrest of reverse transcriptase (RT) progression on the rRNA template due to its inability to incorporate a complementary nucleotide into the synthesized cDNA at the position complementary to the methylated adenine. Since nucleotide A2503 in the wild-type (WT) 70S ribosome is already methylated at position C2 by the housekeeping rRNA-methyltransferase RlmN, we expected the cDNA arrest product to be observed even in the case of 23S rRNA isolated from WT *Tth* cells. Therefore, as a reference and negative control, we used 23S rRNA isolated from a Δ*rlmN Tth* HB27 strain lacking A2503-C2-methylation.

By optimizing the concentrations of nucleotides and RT enzymes, as well as RT reaction time, we developed primer extension reaction conditions that allowed us to discriminate between C2,C8-dimethylated (+*cfr*), C2-monomethylated (WT), and unmodified (Δ*rlmN*) adenine nucleotides at position 2503 (**Figure 1C, D**, lanes 2, 3, and 4, respectively). As expected, the strongest RT arrest was observed for the 23S rRNA isolated from the *cfr-*positive *Tth* cells (**Figure 1C, D**, lanes 4), confirming that PfuCFR effects the desired A2503-C8-methylation. By optimizing the growth conditions of the PfuCFR-expressing *Tth* strain, we achieved levels of A2503-C8-methylation as high as 70% (**Figure 1D**). Since X-ray crystallography is an averaging technique, structural data can be confidently interpreted if at least half of the ribosomes are methylated at position A2503. Altogether, our microbiological and biochemical data show that the Cfr homologs from *P. fimeticola* and *P. fulgidum* possess methyltransferase activities and can be expressed in *Tth* HB27 cells to effect C8-methylation of A2503 in the bacterial 23S rRNA.

### Structure of the 70S ribosome with m^2^m^8^A2503

We purified 70S ribosomes from the *Tth* HB27 strain expressing PfuCFR-methyltransferase for structural analysis. To assess if the presence of aa-tRNA substrates affects the position of C8-methylated nucleotide A2503, we used these Cfr-modified *Tth* 70S ribosomes to assemble a complex with Phe-tRNA^Phe^ and fMet-tRNA_i_^Met^ in the A and P sites, respectively. The complex was crystallized using previously published conditions^19,20,24^, and its structure was determined at 2.55-Å resolution (**Table S3**). At this resolution, nucleotide methylations can be directly visualized in the unbiased difference electron density maps (**Figure 2A**, green mesh), allowing for accurate modeling of the C2,C8-dimethylated nucleotide A2503 (m^2^m^8^A2503) of the 23S rRNA in the structure (**Figure 2A, B**). Additionally, by using non-hydrolyzable, amide-linked aa- and peptidyl-tRNAs (see Online Methods), we were able to capture the PTC in its pre-peptide bond formation state (**Figure S3A**). Consistent with the recent structure of a Cfr-modified *Escherichia coli* 50S ribosomal subunit by the Fujimori group (**Figure S4A, C**)^17^, our structure reveals that C8-methylation does not affect the overall position of nucleotide A2503 in the Cfr-modified ribosome (**Figure S3B**).

**Figure 2.**
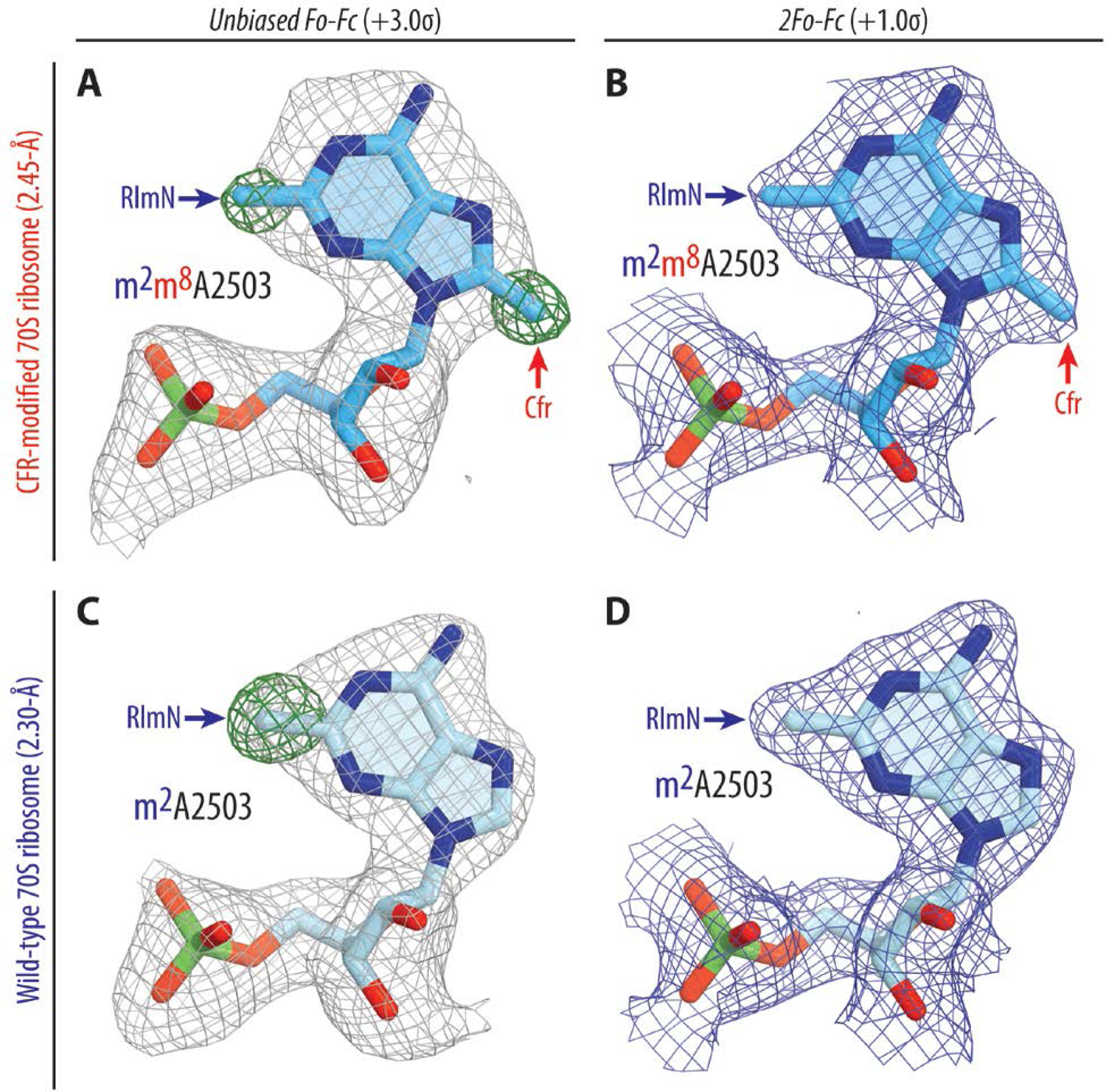
Electron density maps of C2,C8-dimethylated (top), and C2-monomethylated (bottom) A2503 residue of the 23S rRNA in *T. thermophilus* 70S ribosome. (**A, C**) Unbiased *F_o_-F_c_* (grey and green mesh) and (**B, D**) 2*F_o_-F_c_* (blue mesh) electron difference Fourier maps of A2503 residue in the *T. thermophilus* 70S ribosome contoured at 3.0σ and 1.0σ, respectively. Grey mesh shows the *F*_o_-*F*_c_ map after refinement with the entire modified nucleotide omitted. Green mesh, reflecting the presence of methyl groups, shows the *F*_o_-*F*_c_ electron density map after refinement with the nucleotide A2503 built as a regular unmethylated adenine. The refined models of Cfr-modified C2,C8-dimethylated (A, B), or regular RlmN-modified C2-monomethylated (C, D) A2503 nucleotide are displayed in the corresponding electron density maps. The structure and the electron density maps of the wild-type ribosome complex (C, D) are from PDB entry 6XHW^20^. Carbon atoms are colored blue for the Cfr-modified A2503 and light blue for the C8-unmethylated A2503; nitrogens are dark blue; oxygens are red, and phosphorus atoms are green.

To address the possibility of Cfr-induced structural rearrangements around the PTC that can result in multidrug resistance, we compared our structure of the 70S ribosome containing m^2^m^8^A2503 with a structure of the 70S ribosome containing WT m^2^A2503^20^. The alignment revealed no significant changes in the positions of the A- and P-site tRNA substrates (**Figure S3A**) or the key functional nucleotides of the 23S rRNA around the PTC (**Figure S3B**). However, we observed an unanticipated conformational change of nucleotide A2062, which now lacks the characteristic rotation and subsequent Hoogsteen base-pairing with m^2^m^8^A2503 in our structure of the Cfr-modified ribosome (**Figure S3B**). In a previously published WT ribosome structure containing aminoacylated Phe-tRNA^Phe^ and fMet-tRNA_i_^Met^ in the A and P sites, respectively, nucleotide A2062 appears in its rotated conformation (**Figure S5A, S6**) and forms direct van der Waals interactions with the formyl-methionyl moiety of the P-site tRNA^20^. However, in our structure of the Cfr-modified ribosome, the electron density corresponding to A2062 appeared weak and fragmented, indicating the variability of its position (**Figure S5B**). To confirm whether the inability of A2062 to rotate and form a Hoogsteen base-pair with m^2^m^8^A2503 is a direct consequence of Cfr-mediated methylation, we sought to solve the structure of a Cfr-methylated 70S ribosome with peptidyl-tRNA in place of fMet-aminoacyl-tRNA_i_^Met^ in the P site. This is because, in our recently published structures featuring peptidyl-tRNAs or their short analogs in the P site^21,25,26^, nucleotide A2062 always appears in the rotated conformation due to a strong H-bond with the amide of the penultimate amino acid residue of the peptidyl-tRNA (**Figure S7A, B**). If A2062 remains unrotated in the Cfr-methylated ribosome despite the possibility of stabilizing interactions with the P-site peptidyl-tRNA, it would support our hypothesis that Cfr methylation precludes the Hoogsteen base-pairing of A2062 with m^2^m^8^A2503.

### Cfr-mediated modification of A2503 prevents A2062 rearrangement

To this end, we used our recently developed chemoenzymatic approach based on native chemical ligation^26^ to produce a non-hydrolyzable peptidyl-tRNA carrying a formyl-Met-Thr-His-Ser-Met-Arg-Cys (fMTHSMRC) heptapeptide moiety. In a previously published structure of the same peptidyl-tRNA in complex with WT 70S ribosome^26^, the rotated conformation of nucleotide A2062 was stabilized not only by direct H-bonding with the fMTHSMRC-peptide moiety (**Figure 3E; Figure S6A**), but also by π-π stacking with the imidazole side chain of the His3 residue (**Figure S7B**), thus further reinforcing the rotated conformation of A2062. Using this peptidyl-tRNA as the P-site substrate, we assembled a complex of Cfr-modified *Tth* 70S ribosome containing Phe-tRNA^Phe^ in the A site and solved its structure at 2.45-Å resolution (**Figure 3A-D; Table S3**). Similar to the structure with P-site fMet-tRNA_i_^Met^, we observed no significant changes in the positions of the A- or P-site tRNA bodies (**Figure S3C**). Unlike the structure with P-site fMet-tRNA_i_^Met^, the electron density corresponding to A2062 in the structure with P-site peptidyl-tRNA is well-defined (**Figure 3D; Figure S5C**), allowing us to determine its position unequivocally. Remarkably, despite potential stabilizing interactions, nucleotide A2062 remains in the unrotated conformation (**Figure 3D-F; Figure S3D, S4B**), suggesting that Cfr-mediated methylation of m^2^A2503 indeed renders it unable to form a symmetrical Hoogsteen-edge base-pair with A2062. No other structural rearrangements are visible in the PTC, except for a shift in the peptide chain (**Figure S3C**). However, this peptide relocation is likely to be the consequence of the unrotated position of A2062 (rather than its cause) because its His3 residue follows and retains π-π stacking with the A2062 nucleobase in the Cfr-modified ribosome (**Figure S7D**).

**Figure 3.**
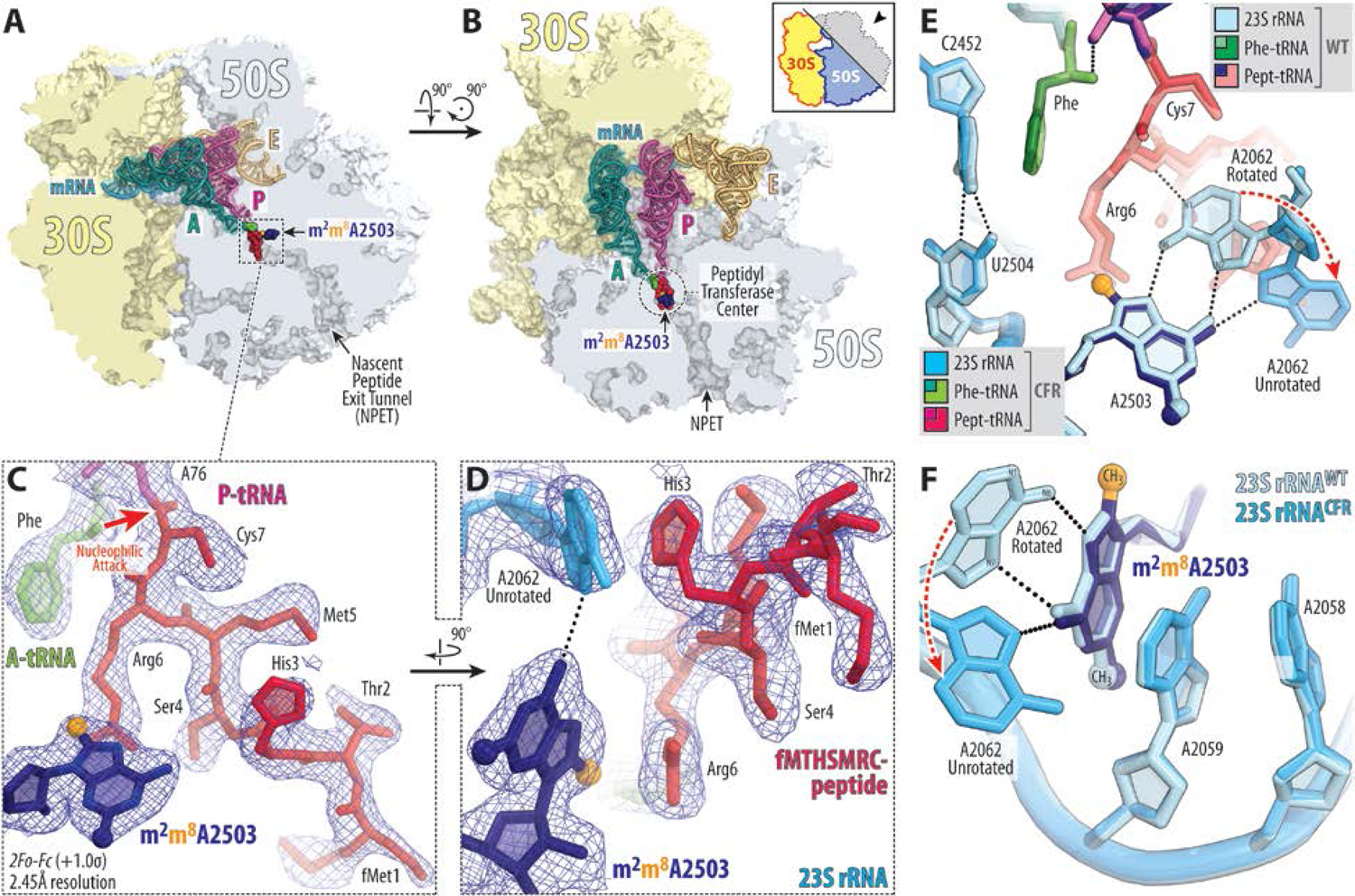
Structure of the Cfr-modified 70S ribosome. (**A, B**) Location of the Cfr-modified nucleotide A2503 (navy blue) carrying the C8-methyl group (orange) in the peptidyl transferase center (PTC) of the *Tth* 70S ribosome relative to tRNAs viewed as cross-cut sections through the nascent peptide exit tunnel (NPET). The 30S subunit is shown in light yellow, the 50S subunit is in light blue, the mRNA is in blue, and the A-, P-, and E-site tRNAs are colored teal, magenta, and light orange, respectively. The phenylalanyl and fMTHSMRC-peptide moieties of the A- and P-site tRNAs are colored green and crimson, respectively. (**C, D**) Close-up views of the 2*F_o_-F_c_* electron density map (blue mesh) of A2503 nucleotide and ribosome-bound A-site Phe-tRNA^Phe^ (teal with amino acid moiety highlighted in green) and P-site fMTHSMRC-peptidyl-tRNA_i_^Met^ (magenta with peptide moiety highlighted in crimson). (**E, F**) Superpositioning of the previously reported structure of wild-type (WT) 70S ribosome containing A-site Phe-tRNA^Phe^ and P-site fMTHSMRC-peptidyl tRNA_i_^Met^ (PDB entry 8CVL^26^) with the structure of the same complexes containing Cfr-methylated nucleotide A2503 in the 23S rRNA. Nucleotides of the Cfr-modified and unmodified ribosomes are shown in blue and light blue, respectively. All structures were aligned based on domain V of the 23S rRNA. E. coli nucleotide numbering is used throughout. H-bonds are shown with dotted lines. Note that the Cfr-modification of nucleotide A2503 does not cause structural rearrangements of the 23S nucleotides around the PTC.

We hypothesize that the observed inability of A2062 to adopt a rotated conformation in the Cfr-modified ribosome leads to resistance against some PTC-acting antibiotics. Indeed, the rotation of A2062 and formation of a Hoogsteen base-pair with m^2^A2503 is required for chloramphenicol^21,25,27^ and its derivatives^28,29^, oxazolidinones^30^, hygromycin A^7^, and A201A^7^ to establish additional binding contacts with the 23S rRNA. An inability to establish this direct contact with the 23S rRNA due to the unrotated conformation of nucleotide A2062 in Cfr-modified ribosomes may thus lead to decreased antibiotic affinity and result in drug resistance. This hypothesis is further supported by the discovery of A2062 spontaneous mutations conferring resistance to chloramphenicol^31^. Further structural analysis show that superpositions of the structure of Cfr-modified *Tth* ribosome with those of the WT ribosome bound to chloramphenicol (**Figure 4A**), linezolid (**Figure 4B**), or hygromycin A (**Figure 4C**) reveal only minor overlaps between the A2503-C8-methyl group and the drug molecules. Therefore, loss of stabilizing interactions between a drug and A2062 in the A2503-C8-methylated ribosome could explain Cfr-mediated resistance towards phenicols, oxazolidinones, hygromycin A, and A201A.

**Figure 4.**
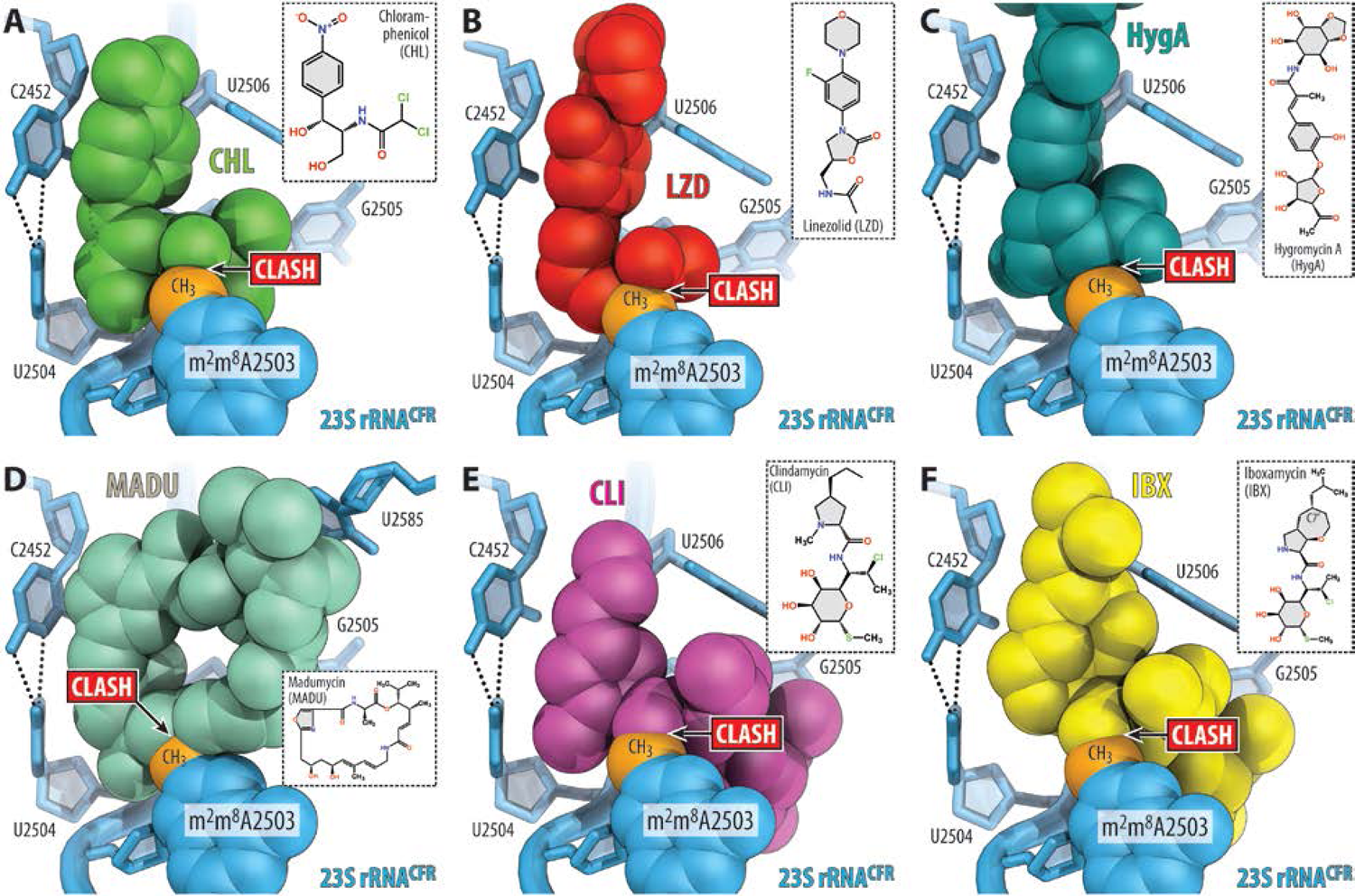
Proposed structural basis for Cfr-mediated resistance to PTC-acting antibiotics. (**A-F**) Superposition of the structures of the Cfr-modified *T. thermophilus* 70S ribosome containing C2,C8-dimethylated A2503 residue in the 23S rRNA (blue) with structures of (wild-type) ribosome-bound antibiotics that target the PTC: chloramphenicol (A, CHL, green, PDB entry 7RQE^25^), linezolid (B, LZD, red, PDB entry 7S1G^30^), hygromycin A (C, HygA, teal, PDB entry 5DOY^7^), madumycin (D, MADU, light teal, PDB entry 5VP2^34^), clindamycin (E, CLI, magenta, PDB entry 4V7V^32^), and iboxamycin (F, IBX, yellow, PDB entry 7RQ8^33^). Note that the C8-methyl group of m^2^m^8^A2503 (highlighted in orange) physically interferes with binding of chemically unrelated antibiotic classes.

In contrast, the rotated conformation of A2062 is not required for binding of lincosamides^32,33^, streptogramins A^34^, pleuromutilins^35^, or 16-membered macrolides^36^. With these drugs, the A2503-C8-methyl group exhibits substantial steric overlaps (**Figure 4D-F**) and, therefore, physically interferes with antibiotic binding. Altogether, our structural analysis shows that the actual mechanism of Cfr-mediated resistance at a molecular level appears to be idiosyncratic for each class of PTC-targeting antibiotics, operating via either an allosteric mechanism (phenicols, oxazolidinones) or direct steric hindrance (lincosamides, streptogramins A, 16-membered macrolides).

### Mechanisms of evading Cfr-mediated resistance

Although the C8-methylation of A2503 leads to high levels of resistance against a range of PTC-targeting antibiotics, including lincosamides, some continue to exhibit activity against Cfr-positive pathogens. The recently disclosed oxepanoprolinamide antibiotic iboxamycin (IBX) exhibits a broad spectrum of activity in high-priority Gram-positive pathogens^33^. Importantly, IBX is highly active against strains harboring Erm-, ABCF-, or Cfr-resistance determinants^33^, although less so against the synergetic protection via 23S modification by Erm/Cfr and direct protection by ABCFs^37,38^. Upon superposition of the Cfr-modified ribosome with that of the ribosome-bound IBX and clindamycin, both molecules are anticipated to exhibit a similar steric clash between the C8-methyl of m^2^m^8^A2503 and the amide moiety of each antibiotic (**Figure 4E, F**). However, *in vitro* susceptibility testing of IBX against *cfr*-expressing strains clearly demonstrate a substantial increase in activity when compared to clindamycin^33^. Simple structural comparison between the antibiotics cannot readily rationalize this increased activity, as the amide moiety remains unchanged between IBX and clindamycin (**Figure 4E, F**). Therefore, a deeper understanding of how IBX maintains interactions with the A2503-C8-methylated ribosome is critical for informing the development of antibiotics with enhanced activity against Cfr-positive pathogens.

To uncover the mechanism of evading Cfr-mediated resistance by IBX, we determined its structure in complex with the Cfr-modified ribosome at 2.55-Å resolution (**Figure 5A, B; Figure S8; Table S3**). Like clindamycin, a network of H-bonds anchors the aminooctose moiety of IBX to nucleotides A2058, A2059, and A2503 in the NPET (**Figure 5C**). However, unlike traditional lincosamides, the bicyclic framework of IBX projects an isobutyl moiety from C7’, conferring additional engagement of the PTC by extending deep into the A-site cleft^33^. This hydrophobic cleft, formed by the 23S rRNA residues A2451 and C2452, normally accommodates the side-chains of incoming amino acids and plays a key role in positioning the aminoacylated 3′-end of A-site tRNA within the PTC during transpeptidation^19,26^. Remarkably, the observed electron-density map reveals that the binding site of IBX in the Cfr-methylated ribosome is nearly identical to that in the WT ribosome (**Figure 5C, D**), whereas m^2^m^8^A2503 undergoes a movement of ∼1–1.5 Å relative to its canonical position in order to accommodate the antibiotic (**Figure 5D**). This previously unknown displacement of m^2^m^8^A2503, which also disrupts one H-bond typically formed between 2’-OH of A2503 and the aminooctose group of lincosamides (**Figure 5D**), is not a trivial concession to make as it is clearly sufficient to disrupt clindamycin binding. However, the extended hydrophobic interaction between IBX and the A-site cleft compensates for this clash with the modified A2503 nucleobase, providing sufficient affinity to evade Cfr-based resistance. Interestingly, this IBX-induced displacement of m^2^m^8^A2503 observed in the Cfr-methylated ribosome is conceptually reminiscent of the previously reported IBX-induced displacement of m^6^A2058 for the Erm-methylated ribosome^33^. This unanticipated nucleotide mobility further underscores a synthetic antibiotic’s ability to overcome methyltransferase-mediated resistance by compensating with binding affinity from proximal interactions, going against conventional small-molecule inhibitor design principles.

**Figure 5.**
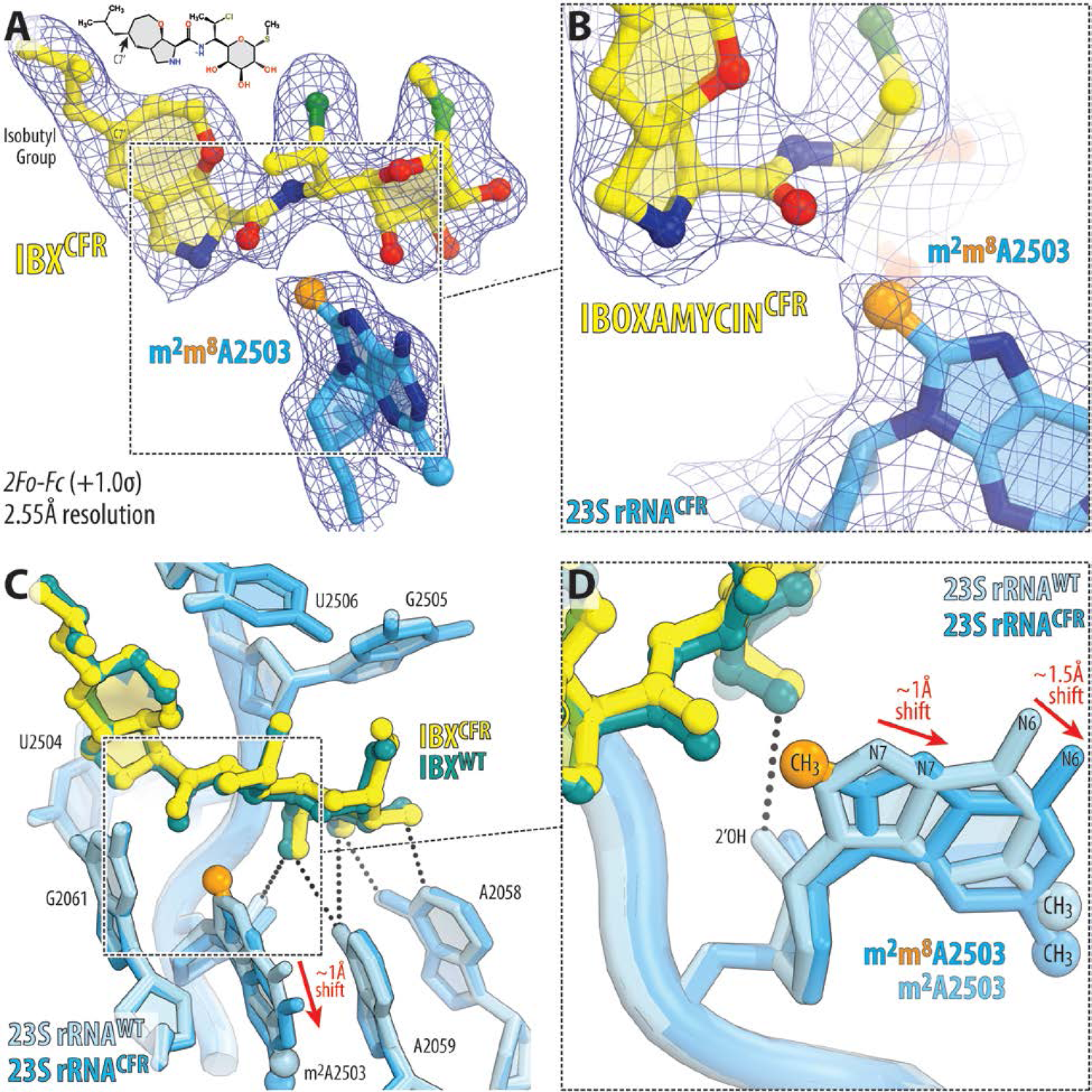
Structure of iboxamycin (IBX) bound to the Cfr-methylated 70S ribosome. (**A, B**) Electron density map (blue mesh) contoured at 1.0σ of IBX (yellow) in complex with the Cfr-modified *T. thermophilus* 70S ribosome containing C2,C8-dimethylated A2503 residue in the 23S rRNA (blue). The C8-methyl group of m^2^m^8^A2503 is highlighted in orange. (**C, D**) Superposition of IBX (teal) in complex with the WT 70S ribosome containing a C8-unmodified residue m^2^A2503 (light blue) and the structure of IBX (yellow) in complex with the Cfr-modified 70S ribosome containing an m^2^m^8^A2503 residue (blue). Hydrogen bonds are depicted with dotted lines. The position of IBX is almost identical in the two structures, while the m^2^m^8^A2503 residue in the IBX-bound Cfr-modified structure is shifted by ∼1–1.5 Å relative to the canonical position of the m^2^m^8^A2503 residue of the wild-type ribosome with IBX bound (red arrows).

Besides PhLOPS_A_ antibiotics, expression of Cfr also confers resistance towards 16-membered macrolides (16MM), which bind in the canonical macrolide binding pocket located in the NPET^36^. However, unlike conventional 14-membered macrolides (such as ERY), these compounds protrude further into the PTC due to their long oligosaccharide extensions (such as in tylosin), with some even encroaching upon the A-site cleft due to further substitutions on the oligosaccharide (such as in josamycin)^36^. While Cfr confers strong (up to 64-fold) resistance to 16MMs such as spiramycin or josamycin^6^, it confers only a 2-fold increase in MIC for tylosin (TYL)^5,6^, suggesting that this drug exhibits at least some binding affinity for the Cfr-modified ribosome. To assess this experimentally, we determined the structure of TYL in complex with Cfr-modified ribosomes, with Phe-tRNA^Phe^ and fMet-tRNA_i_^Met^ in the A and P sites, respectively. The 2.65-Å electron density map revealed TYL bound in the canonical macrolide NPET binding pocket, together with the Cfr-modified m^2^m^8^A2503 nucleotide (**Figure 6A; Table S3**). Consistent with the previous 3-Å structure of TYL in complex with an A2503-unmodified, ligand-free 50S subunit from the archaeon *Haloarcula marismortui*^36^, the acetaldehyde motif at C6 of TYL forms a covalent bond with the exocyclic N6-amino group of A2062 in the *Tth* ribosome, as manifested by the continuous electron density connecting the drug to the nucleobase (**Figure 6B**). To accurately evaluate the effect of Cfr-methylation on the position of TYL in the Cfr-modified ribosome, we also determined its structure in complex with the unmodified WT *Tth* 70S ribosome (**Figure 6C; Table S3**).

**Figure 6.**
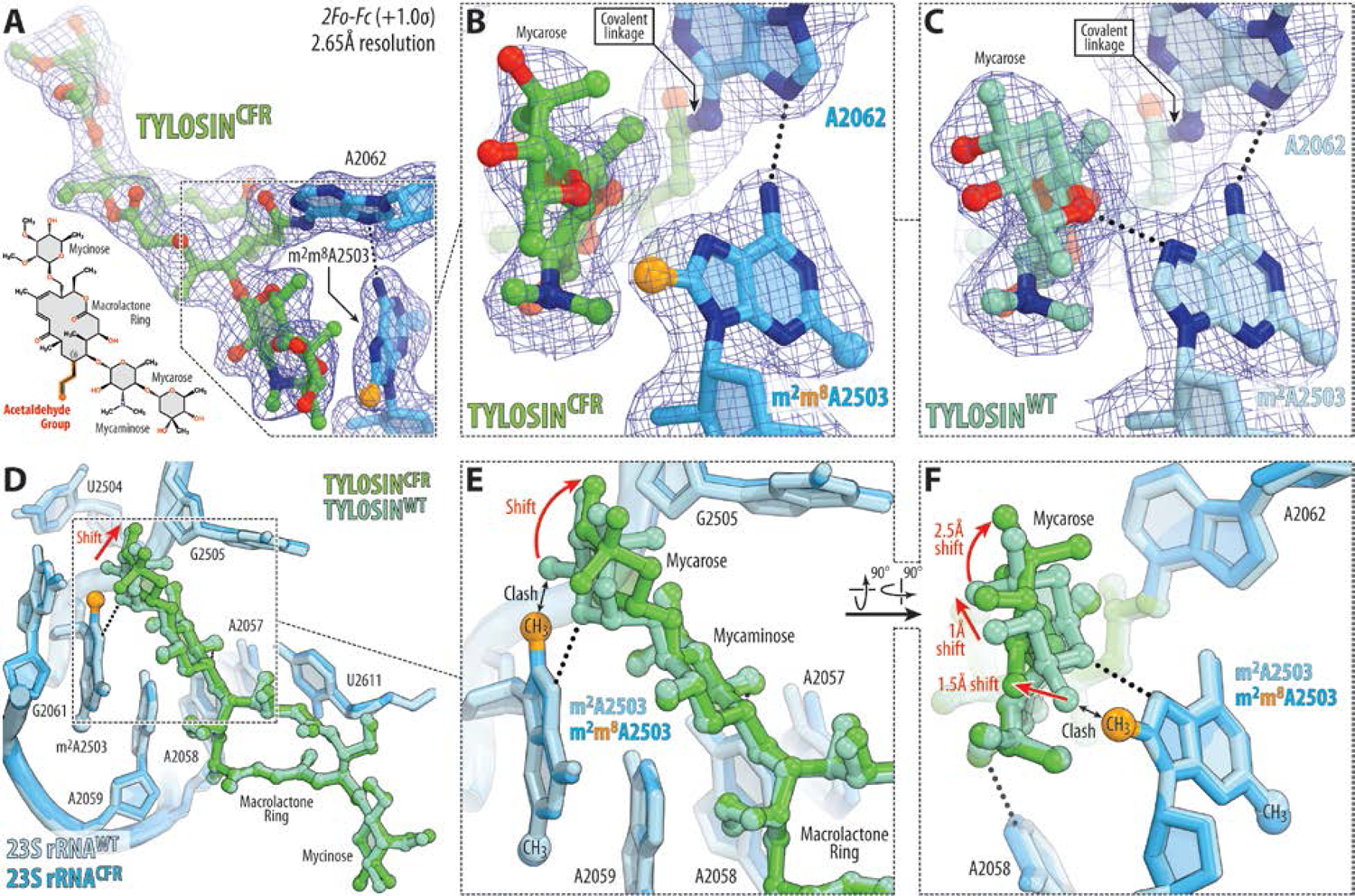
Structure of tylosin (TYL) bound to the Cfr-modified and WT 70S ribosomes. (**A-C**) Electron density map (blue mesh) contoured at 1.0σ of TYL (green) in complex with the Cfr-modified (A, B) or wild-type (C) *T. thermophilus* 70S ribosome containing m^2^m^8^A2503 (blue) or m^2^A2503 (light blue) residues in the 23S rRNA, respectively. The C8-methyl group of m^2^m^8^A2503 is highlighted in orange. (**D-F**) Superposition of TYL (light teal) in complex with the WT 70S ribosome containing a C8-unmodified residue m^2^A2503 (light blue) and the structure of TYL (green) in complex with the Cfr-modified 70S ribosome containing an m^2^m^8^A2503 residue (blue). Hydrogen bonds are depicted with dotted lines. Note that the position of m^2^m^8^A2503 residue is almost identical in the two structures, whereas TYL binding to the Cfr-modified ribosome causes ∼1.5–2.5 Å shift of its mycarose moiety to a new position (red arrows).

Alignment of our structure of Cfr-modified 70S ribosomes with the previous structures of 16MMs in complex with the *H. marismortui* 50S subunit^36^ reveals a substantial steric clash between the mycarose moiety of 16MMs and the A2503-C8-methyl group (**Figure S9**), thus explaining Cfr-mediated resistance to this subclass of macrolides. Curiously, alignment of our structures of Cfr-modified ribosomes, with and without TYL bound, reveals neither displacement of nucleotide A2503 nor any structural rearrangements elsewhere in the 23S rRNA (**Figure S10**), suggesting that TYL must change its position to avoid clashes with the C8-methyl group of m^2^m^8^A2503. Indeed, a comparison between our structures of ribosome-bound TYL, one in complex with WT ribosome and the other with the Cfr-modified ribosome, shows that while the overall drug binding site remains the same in both structures, the mycarose moiety of TYL in the Cfr-modified ribosome moves away from m^2^m^8^A2503 to avoid clashing with its C8-methyl group (**Figure 6D–F**). This observation is in striking contrast to IBX, which instead displaces m^2^m^8^A2503 from its canonical position (**Figure 5C, D**). Thus, these examples suggest that drugs active against Cfr-expressing bacteria can employ distinct mechanisms to evade Cfr-based resistance: (i) strong target engagement resulting in displacement of the Cfr-methylated m^2^m^8^A2503 nucleotide, as in the case of IBX; or (ii) flexible target engagement allowing the drug to avoid clashes with the m^2^m^8^A2503 nucleotide, as in the case of TYL.

## CONCLUSIONS

Altogether, our structural study of Cfr-mediated resistance to PTC-targeting ribosomal antibiotics has uncovered at least two distinct molecular mechanisms: (i) direct steric hindrance of the A2503-C8-methyl group with the ribosome-bound drug confers resistance to lincosamides and streptogramins A; and (ii) Cfr-methylation-induced rearrangement of the nucleotide A2062 to a conformation incompatible with drug binding confers resistance to phenicols, oxazolidinones, and hygromycin A. The structure of iboxamycin in complex with the Cfr-modified ribosome revealed an unexpected finding: by establishing strong contacts with the 23S rRNA, iboxamycin is able to displace m^2^m^8^A2503 from its canonical position, thereby evading Cfr-based resistance. Thus, forming new interactions with the ribosomal A site may prove a general strategy in the design of expanded-spectrum antibiotics capable of overcoming both Cfr- and Erm-methyltransferase-mediated resistance. We believe that the information provided by the structure of the Cfr-modified ribosome and its complexes with antibiotics is an indispensable starting point for the structure-based development of next-generation drugs active against the most challenging multidrug-resistant pathogens. This quest will likely be stimulated by the recently discovered combinatorial approaches for the synthesis of novel oxepanoprolinamides^33^, streptogramins^39^, pleuromutilins^40^, and other antibiotics.

## ACKNOWLEDGMENTS

We thank the staff at NE-CAT beamlines 24ID-C and 24ID-E for help with X-ray diffraction data collection, especially Drs. Malcolm Capel, Frank Murphy, Surajit Banerjee, Igor Kourinov, David Neau, Jonathan Schuermann, Narayanasami Sukumar, Anthony Lynch, James Withrow, Kay Perry, Ali Kaya, and Cyndi Salbego.

This work is based upon research conducted at the Northeastern Collaborative Access Team beamlines, which are funded by the National Institute of General Medical Sciences from the National Institutes of Health [P30-GM124165 to NE-CAT]. The Eiger 16M detector on 24-ID-E beamline is funded by an NIH-ORIP HEI grant [S10-OD021527 to NE-CAT]. This research used resources of the Advanced Photon Source, a US Department of Energy (DOE) Office of Science User Facility operated for the DOE Office of Science by Argonne National Laboratory under Contract No. DE-AC02-06CH11357. This work was supported by the National Institute of Allergy and Infectious Diseases of the National Institutes of Health [R01-AI168228 to A.G.M. and R21-AI163466 to Y.S.P.], National Institute of General Medical Sciences of the National Institutes of Health [R01-GM094157 to S.T.G. and R01-GM132302 to Y.S.P.], USDA National Insititute for Food and Agriculture [Hatch Project 1016013 to S.T.G.], the Illinois State startup funds [to Y.S.P.], the Swedish Research Council (Ventenskapsrådet) [2019-01085 and 2022-01603 to G.C.A.], the Knut and Alice Wallenberg Foundation [2020.0037 to G.C.A.], and the Carl Tryggers Stiftelse för Vetenskaplig Forskning [CTS19:24 to G.C.A.]. K.J.Y.W. was supported by a National Science Scholarship (Ph.D.) from the Agency for Science, Technology and Research (Singapore). The funders had no role in study design, data collection and analysis, decision to publish, or manuscript preparation.

## AUTHOR CONTRIBUTIONS

E.V.A. with the help from S.M.B. and M.S.S. constructed the *T. thermophilus* HB27 strain expressing Cfr-like methyltransferases; G.C.A. performed phylogenetic analysis and identified putative thermostable *cfr*-like genes; B.I.C.T. and K.J.Y.W. synthesized iboxamycin; E.E.K. prepared Δ*rlmN* knock-out *T. thermophilus* HB27 strain; E.V.A. performed the assessment of A2503-C8-methylation using primer extension assay; E.V.A. and E.A.S. grew *T. thermophilus* cells and purified Cfr-modified 70S ribosomes; E.A.S. and E.V.A. prepared hydrolysis-resistant aminoacyl- and peptidyl-tRNAs; E.V.A., E.A.S., and Y.S.P. designed and performed X-ray crystallography experiments; A.G.M., Y.S.P., and S.T.G. supervised the experiments. All authors interpreted the results. E.V.A., B.I.C.T., K.J.Y.W, A.G.M., and Y.S.P. wrote the manuscript.

## COMPETING INTERESTS STATEMENT

A.G.M. is an inventor in a provisional patent application submitted by the President and Fellows of Harvard College covering oxepanoprolinamide antibiotics described in this work. A.G.M. has filed the following international patent applications: WO/2019/032936 ‘Lincosamide Antibiotics and Uses Thereof’ and WO/2019/032956 ‘Lincosamide Antibiotics and Uses Thereof’. All other authors declare no competing financial or non-financial interests.

## ONLINE METHODS

### Reagents

Unless stated otherwise, all chemicals and reagents were obtained from MilliporeSigma (USA). Iboxamycin was synthesized following the protocols described previously^41^. Tylosin was obtained from MilliporeSigma (USA).

### Phylogenetic analysis of cfr genes

To identify a large set of Cfr homologs from which proteins encoded by thermophiles could be selected, a BlastP search against the NCBI (https://www.ncbi.nlm.nih.gov/) Refseq database was carried out, with the CfrA from the Comprehensive Antibiotic Resistance Database (CARD)^42^ as the query. An E value cut-off of 1e^-70^ was used, which allowed all available Cfr sequences to be identified, along with a subset of RlmN family homologs. The sequences were aligned using MAFFT-L-INS-I v6.861b^43^. After removing alignment positions with more than 50% gaps with TrimAl v 1.2^44^, preliminary phylogenetic analysis was carried out with FastTree v 2.1^45^ to distinguish Cfr representatives from the more distantly related RlmN, and identify thermophilic Cfr representatives. For the representative tree shown in **Figure S1**, additional Cfr and RlmN representatives downloaded respectively from the CARD^42^ and Uniprot^46^ databases were aligned as above with the thermophile Cfr and other Cfr sequences of interest to this study. After a TrimAl step as above, phylogenetic analysis was carried out with IQTree version 2.1.2^47^ on the CIPRES Science Gateway^48^ with 1000 rapid bootstrap replicates and automatic model determination. All alignments and phylogenies are available from https://github.com/GCA-VH-lab/2023_Cfr.

### Construction of the Cfr(+) Thermus thermophilus HB27 strain

The original *cfr* gene from mesophilic *Staphylococcus aureus* (*Sau*) encoding for A2503-C8-methyltransferase as well as several *cfr*-homologous genes from various moderately thermophilic bacterial species (**Supplementary Table 1**) were commercially synthesized *de novo* (the synthesis was carried out by GenScript, Inc.) and cloned into the pBGAA1-BthERM expression vector, which we have generated in our previous study^20^. The DNA sequences of the synthesized genes were adjusted to the codon usage for optimal gene expression in the *Thermus thermophilus* (*Tth*) HB27 host. The synthesized *cfr*-like genes were inserted in place of the *erm* gene, using *Nsi*I and *Ssp*I unique restriction sites, and placed under the control of the inducible Parg promoter (Figure 1B, orange). This vector originates from the parent pBGAA1 plasmid^49^, which was specifically designed to replicate in *Tth* due to the presence of repA gene and oriV2 replication origin (Figure 1B). It also carries the *hph* gene^50^ that provides a high level of resistance to hygromycin B (HygB) at a broad range of temperatures from 37°C to 65°C and allows for positive selection (Figure 1B, blue). The resulting *cfr*-expression vectors (Figure 1B) were transformed into the *Tth* HB27 host, and cells were then propagated in standard liquid ATCC 697 medium supplemented with Castenholz salt. Transformants were plated on 3% agar plates prepared with the same medium. The agar plates were incubated at 60°C for 48 hours to allow colony formation. To select the Cfr(+) colonies, transformed *Tth* cells were plated on agar containing 50 µg/ml HygB. Individual HygB-resistant colonies were picked, diluted in a fresh medium containing 50 µg/ml HygB and grown at 58°C.

### Construction of ΔrlmN Thermus thermophilus HB27 strain

Upstream and downstream homology regions (UHR and DHR, respectively) flanking the *Tth rlmN* coding sequence were amplified using the following primer pairs:

- 5’-TAAAACGACGGCCAGTGCCACCTCGAGGCCCTTCGCCC-3’ (UHR-Fwd);
- 5’-GTCCTTTCATACCCTCCCATTGTAGCCGAGAAG-3’ (UHR-Rev);
- 5’-ACCATTTTGATGACCCTCACACCGCTTCCAGAAG-3’ (DHR-Fwd);
- 5’-AGTCGACCTGCAGGCATGCACGCCTCGGACACGGCGCA-3’ (DHR-Rev).

The *htk* gene encoding a thermostable kanamycin adenyltransferase^51^ was amplified using primer pair:

- 5’-ATGGGAGGGTATGAAAGGACCAATAATAATGAC-3’ (HTK-Fwd);
- 5’-GTGAGGGTCATCAAAATGGTATGCGTTTTG-3’ (HTK-Rev).

The obtained UHR, *htk*, and DHR sequences were inserted into *HindIII*-digested plasmid pUC18 using the NEBuilder HiFi DNA Assembly Master Mix (E2621, New England Biolabs) following the manufacturer’s protocol. The resulting assembly was used to transform NEB 5-alpha competent *E. coli* cells (C2987, New England Biolabs), and transformants were selected on LB ampicillin plates. *Tth* strain HB27 was transformed with the resulting plasmid, and recombinants were selected on TEM kanamycin plates. Individual *Tth* isolates were purified, sequencednto confirm the replacement of *rlmN* gene with *htk* gene, and also analyzed by diagnostic PCR using primer pair #1:

- 5’-ATTCGACATATGGCCGCTTCCCACGCCCTC-3’;
- 5’-TTTTTCATATGATACCTCCTGTCATCGCCCGGCGCC-3’;

and primer pair #2:

- 5’-ATGAAAGGACCAATAATAATGACTAGAGAAGAAAGAATG-3’;
- 5’-TCAAAATGGTATGCGTTTTGACACATCCACTATATATCC-3’

### Assessment of Cfr expression

Minimal inhibitory concentrations (MICs) of various PTC-targeting drugs, such as chloramphenicol (CHL), florfenicol (FFL), lincomycin (LNC), clindamycin (CLI), or iboxamycin (IBX) as well as NPET-acting macrolide ERY against constructed Cfr(+) *T. thermophilus* HB27 strains were determined as described previously^52^. Untransformed parent or transformed with the empty vector strains were used as negative controls (**Table S2**, values highlighted in blue). To minimize the negative effect of high temperature on the structure and activity of Cfr-methyltransferases, the MIC testing experiments were performed at 55-58°C.

### Isolation and purification of A2503-C2,C8-dimethylated 70S ribosomes

The *Tth* HB27 cells expressing Cfr-like methyltransferase from *P. fulgidum* were grown at 58-60°C in flasks with a total of 9 liters of ATCC 697 medium with Castenholz salt and also supplemented with 50 µg/ml HygB (to retain the Cfr expression vector) and 50 µg/ml CLI (to sustain high levels of induction of PfuCFR gene expression). The Cfr(+) *Tth* cells were harvested at early-to-mid log-phase (OD_600_ = 0.9-1.0) and used for the subsequent large-scale preparation of A2503-C2,C8-dimethylated 70S ribosomes. The total yield was approximately 19 grams of cell paste. Purification of Cfr-methylated ribosomes was accomplished as optimized previously for the A2058-N6-dimethylated^20,33^ as well as the wild-type^19,24^ 70S ribosomes from *T. thermophilus*. We routinely use this procedure for the preparation of ribosomes for our crystallographic studies. The main steps of ribosome purification included cell lysis, sucrose cushion ultracentrifugation, reverse-phase chromatography, and, finally, separation of the tightly-coupled 70S ribosomes from individual subunits by sucrose gradient centrifugation. The final 70S pellets were suspended in a buffer, flash-frozen in liquid nitrogen, and stored at −80°C until used in crystallization experiments.

### Primer extension analysis

To assess the extent of A2503-C8-methylation, 23S rRNA isolated from the Cfr(+), wild-type or *ΔrlmN T. thermophilus* HB27 cells was analyzed by primer extension with SuperScript III Reverse Transcriptase (Invitrogen, USA) according to the manufacturer’s protocol. To this end, 2 pmol of radioactively 5′-[^32^P]-labeled primer (5′-GCCCGTGGCGGATAGAGACCG-3′ or 5′-TCTTCAGCCCCAGGATGCGACGAGCCG-3′) was annealed with 1.25 µg of each of the three 23S rRNA samples. The primer extension reactions by reverse transcriptase were carried out at 50°C in the presence of 0.5 mM of each dNTP (Figure 1C) or 1 mM dATP, dCTP, dTTP, and 0.2 mM ddGTP (Figure 1D). For the reactions with ddGTP, the extension time was decreased to 5 minutes, and the concentration of reverse transcriptase was reduced 4-fold relative to the suggested in the manufacturer’s protocol. Primer extension cDNA products were purified by phenol extraction, precipitated with ethanol, and resolved on 6% polyacrylamide sequencing gels in TBE buffer. Gels were transferred onto Whatman paper, dried, exposed to the phosphorimager screen overnight, and visualized in a Typhoon RGB phosphorimager (Cytiva, USA).

### X-ray crystallographic structure determination

To obtain high-resolution structures that would allow visualizing of structural features crucial for this study, we employed a strategy that relies on the use of hydrolysis-resistant aminoacylated tRNAs, which exhibit higher (than deacylated tRNAs) affinity to the 70S ribosome and better stabilize it in the unrotated state resulting in noticeably higher resolution of the resulting datasets. Moreover, such complexes represent the functional pre-attack states of the ribosome. Synthetic mRNA with the sequence 5’-GGC-AAG-GAG-GUA-AAA-**AUG**-**UUC**-UAA-3’ containing Shine-Dalgarno sequence followed by the P-site methionine and the A-site phenylalanine codons was obtained from Integrated DNA Technologies (USA). Non-hydrolyzable aminoacylated tRNAs, Phe-NH-tRNA^Phe^ and fMet-NH-tRNA_i_^Met^, were prepared as described previously^19,53^.

Complexes of the A2503-C2,C8-dimethylated *T. thermophilus* 70S ribosomes with mRNA and hydrolysis-resistant A-site aminoacyl (Phe-tRNA^Phe^) and P-site aminoacyl (fMet-tRNA_i_^Met^) or peptidyl (fMTHSMRC-tRNA_i_^Met^) tRNAs were formed as described previously for deacylated ^24^ or aminoacylated tRNAs^19,20^. For *Tth* 70S ribosome complexes with IBX or TYL, the antibiotic was included in the stabilization buffers (50 µM of IBX and 1 mM of TYL).

Collection and processing of the X-ray diffraction data, model building, and structure refinement were performed as described in our previous reports^19–21,24,26,33^. Diffraction data were collected at beamlines 24ID-C and 24ID-E at the Advanced Photon Source (Argonne National Laboratory). A complete dataset for each complex was collected using 0.979 Å irradiation at 100 K from multiple regions of the same crystal, using 0.3-degree oscillations. Raw data were integrated and scaled using XDS software (Jan 10, 2022)^54^. Molecular replacement was performed using PHASER from the CCP4 program suite (version 7.0)^55^. The search model was generated from the previously published structures of *T. thermophilus* 70S ribosome with bound mRNA and aminoacylated tRNAs (PDB entries 6XHW^20^ or 8CVL^26^). Initial molecular replacement solutions were refined by rigid-body refinement with the ribosome split into multiple domains, followed by positional and individual B-factor refinement using PHENIX software (version 1.17)^56^. Non-crystallographic symmetry restraints were applied to four parts of the 30S ribosomal subunit (head, body, spur, and helix 44) and four parts of the 50S subunit (body, L1-stalk, L10-stalk, and C-terminus of the L9 protein). Structural models were built in Coot (version 0.8.2)^57^. Structural models and restraints for IBX and TYL were generated using PRODRG online software (http://prodrg1.dyndns.org)^58^. The statistics of data collection and refinement are compiled in **Table S3**. All figures showing atomic models were rendered using PyMol software (www.pymol.org).

## DATA AVAILABILITY STATEMENT

Coordinates and structure factors were deposited in the RCSB Protein Data Bank with accession codes:

- **8G29** for the A2503-C2,C8-dimethylated *T. thermophilus* 70S ribosome in complex with mRNA, aminoacylated A-site Phe-NH-tRNA^Phe^, aminoacylated P-site fMet-NH-tRNA_i_^Met^, and deacylated E-site tRNA^Phe^;
- **8G2A** for the A2503-C2,C8-dimethylated *T. thermophilus* 70S ribosome in complex with mRNA, aminoacylated A-site Phe-NH-tRNA^Phe^, peptidyl P-site fMTHSMRC-NH-tRNA_i_^Met^, and deacylated E-site tRNA^Phe^.
- **8G2B** for the A2503-C2,C8-dimethylated *T. thermophilus* 70S ribosome in complex with mRNA, deacylated A-site tRNA^Phe^, aminoacylated P-site fMet-NH-tRNA_i_^Met^, deacylated E-site tRNA^Phe^, and iboxamycin;
- **8G2C** for the A2503-C2,C8-dimethylated *T. thermophilus* 70S ribosome in complex with mRNA, aminoacylated A-site Phe-NH-tRNA^Phe^, aminoacylated P-site fMet-NH-tRNA_i_^Met^, deacylated E-site tRNA^Phe^, and tylosin;
- **8G2D** for the wild-type *T. thermophilus* 70S ribosome in complex with mRNA, deacylated A-site tRNA^Phe^, deacylated P-site tRNA_i_^Met^, deacylated E-site tRNA^Phe^, and tylosin;

All previously published structures that were used in this work for structural comparisons were retrieved from the RCSB Protein Data Bank: PDB entries 6XHW, 8CVL, 7LVK, 7RQE, 7S1G, 5DOY, 5VP2, 4V7V, 7RQ8, 1K9M, 1KD1, 1K8A.

No sequence data were generated in this study. Analyzed protein sequences are presented with their corresponding accession numbers in the phylogenetic tree (**Figure S1**) for retrieval from the NCBI protein database.

