## Supplementary Information for "Structural basis of Cfr-mediated antimicrobial resistance and mechanisms for its evasion"

**This file includes:**

I. Supplementary Tables 1 to 3;

II. Supplementary Figures 1 to 9 with legends;

III. Supplementary References.

### I. SUPPLEMENTARY TABLES

**Supplementary Table 1 | Moderately thermophilic bacteria carrying *cfr*-like genes used in this study.** Percent identity of amino acid sequences of Cfr-like proteins from thermophilic bacteria with the sequences of well-characterized Cfr-methyltransferase from a clinical isolate of *S. aureus*.

| Bacterial speicies name | Optimal growth temperature, ºC | Maximum growth temperature, ºC | Protein identity with Cfr, % |
|---|---|---|---|
| *Clostridium sporogenes (Csp)*[1] | **45-50°C** | **55°C** | **57.1%** |
| *Thermosporothrix hazakensis (Tha)*[2] | **50°C** | **58°C** | **58.5%** |
| *Desmospora activa (Dac)*[3] | **30-50°C** | **50°C** | **58.3%** |
| *Planifilum fimeticola (Pfi)*[3,4] | **55-63°C** | **65°C** | **57.7%** |
| *Planifilum fulgidum (Pfu)*[3,4] | **60-65°C** | **67°C** | **57.7%** |

**Supplementary Table 2 | Minimal inhibitory concentrations (MICs) of various antibiotics against *Thermus thermophilus* HB27 cells transformed with pBGAA1 vectors carrying indicated *cfr*-like genes.** CHL – chloramphenicol, FFL – florfenicol, LNC – lincomycin, CLI – clindamycin, IBX – iboxamycin, ERY – erythromycin, N/D – not determined.

| Transformed vector | MIC, µg/ml (fold change relative to WT) | | | | | |
|---|---|---|---|---|---|---|
| | CHL | FFL | LNC | CLI | IBX | ERY |
| *Tth* HB27 (WT parent strain) | 2 | 0.5 | 2 | 0.125 | 0.125 | 8 |
| pBGAA1 (empty vector) | 2 | 0.5 | 2 | 0.125 | 0.125 | 4 |
| pBGAA1-SauCFR | 2 | N/D | 2 | 0.125 | N/D | 0.5 |
| pBGAA1-CspCFR | 2 | N/D | 2 | 0.125 | N/D | 2 |
| pBGAA1-ThaCFR | 2 | N/D | 4 | 0.125 | N/D | 2 |
| pBGAA1-DacCFR | 2 | N/D | 4 | 0.125 | N/D | 2 |
| pBGAA1-PfiCFR | 64 (32x) | N/D | 2048 (1024x) | >512 (>4096x) | N/D | 2 (0.25x) |
| pBGAA1-PfuCFR | 64 (32x) | 32 (64x) | 2048 (1024x) | >512 (>4096x) | 8 (64x) | 2 (0.25x) |

**Supplementary Table 3 | X-ray data collection and refinement statistics.**

| ***Crystals*** | | A2503-C2,C8-dimethylated 70S ribosome in complex with A-site Phe-tRNA$^{Phe}$, and P-site fMet-tRNA$_i^{Met}$ PDB entry 8G29 | A2503-C2,C8-dimethylated 70S ribosome in complex with A-site Phe-tRNA$^{Phe}$, and P-site fMTHSMRC-tRNA$_i^{Met}$ PDB entry 8G2A | A2503-C2,C8-dimethylated 70S ribosome in complex with A-site tRNA$^{Phe}$, P-site fMet-tRNA$_i^{Met}$, and IBX PDB entry 8G2B |
|---|---|---|---|---|
| ***Diffraction data*** | | | | |
| Space Group | | $P2_12_12_1$ | $P2_12_12_1$ | $P2_12_12_1$ |
| Unit Cell Dimensions, Å (a x b x c) | | 210.09 x 449.74 x 625.05 | 210.01 x 450.38 x 625.75 | 209.64 x 449.56 x 621.12 |
| Wavelength, Å | | 0.97911 | 0.97918 | 0.97918 |
| Resolution range (outer shell), Å | | 182-2.55 (2.62-2.55) | 190-2.45 (2.72-2.45) | 182-2.55 (2.62-2.55) |
| $I/\sigma I$ (outer shell) | | 10.57 (0.97) | 8.76 (0.86) | 8.80 (0.92) |
| Resolution at which $I/\sigma I$=1, Å | | 2.55 | 2.45 | 2.55 |
| Resolution at which $I/\sigma I$=2, Å | | 2.80 | 2.70 | 2.80 |
| CC(1/2) at which $I/\sigma I$=1, % | | 19.4 | 22.8 | 16.6 |
| CC(1/2) at which $I/\sigma I$=2, % | | 50.0 | 55.0 | 50.0 |
| Completeness (outer shell), % | | 99.1 (99.1) | 99.2 (97.7) | 98.2 (98.5) |
| $R_{merge}$ (outer shell)% | | 13.6 (172.8) | 19.2 (248.4) | 15.7 (174.6) |
| No. of crystals used | | 1 | 1 | 1 |
| No. of Reflections Used: | Total | 9,731,340 | 21,563,296 | 9,628,355 |
| | Unique | 1,877,927 | 2,124,013 | 1,845,878 |
| Redundancy (outer shell) | | 5.18 (5.40) | 10.15 (10.28) | 5.22 (5.39) |
| ***Refinement*** | | | | |
| Resolution range of the diffraction data included in the refinement, Å | | 154-2.55 | 125-2.45 | 123-2.55 |
| $R_{work}/R_{free}$, % | | 20.9/25.6 | 20.8/25.2 | 21.3/26.4 |
| *No. of Non-Hydrogen Atoms* | | | | |
| RNA | | 200,309 | 200,289 | 200,203 |
| Protein | | 90,976 | 91,072 | 90,976 |
| Ions (Mg, K, Zn, Fe) | | 2,829 | 2,813 | 2,823 |
| Waters | | 4,358 | 4,363 | 4,350 |
| *Ramachandran Plot* | | | | |
| Favored regions, % | | 89.78 | 91.80 | 90.97 |
| Allowed regions, % | | 8.20 | 7.98 | 8.81 |
| Outliers, % | | 2.02 | 0.22 | 0.22 |
| *Deviations from ideal values (RMSD)* | | | | |
| Bond, Å | | 0.009 | 0.010 | 0.008 |
| Angle, degrees | | 1.468 | 1.409 | 1.407 |
| Chirality | | 0.060 | 0.059 | 0.058 |
| Planarity | | 0.007 | 0.008 | 0.007 |
| Dihedral, degrees | | 17.339 | 17.096 | 17.467 |
| Average B-factor (overall), Å$^2$ | | 60.3 | 64.1 | 62.2 |

**Supplementary Table 3 | X-ray data collection and refinement statistics.**

| ***Crystals*** | | A2503-C2,C8-dimethylated 70S ribosome in complex with A-site Phe-tRNA$^{Phe}$, P-site fMet-tRNA$_i^{Met}$, and Tylosin PDB entry 8G2C | Wild-type 70S ribosome in complex with A-site tRNA$^{Phe}$, P-site tRNA$_i^{Met}$, and Tylosin PDB entry 8G2D |
|---|---|---|---|
| ***Diffraction data*** | | | |
| Space Group | | $P2_12_12_1$ | $P2_12_12_1$ |
| Unit Cell Dimensions, Å (a x b x c) | | 209.65 x 448.12 x 621.73 | 209.36 x 449.77 x 616.87 |
| Wavelength, Å | | 0.97911 | 0.97910 |
| Resolution range (outer shell), Å | | 174-2.65 (2.72-2.65) | 254-2.70 (2.72-2.70) |
| I/σI (outer shell) | | 8.49 (1.02) | 7.32 (0.99) |
| Resolution at which I/σI=1, Å | | 2.65 | 2.70 |
| Resolution at which I/σI=2, Å | | 2.85 | 2.90 |
| CC(1/2) at which I/σI=1, % | | 25.3 | 20.5 |
| CC(1/2) at which I/σI=2, % | | 55.0 | 50.0 |
| Completeness (outer shell), % | | 98.7 (99.6) | 97.2 (94.3) |
| $R_{merge}$ (outer shell)% | | 12.4 (120.3) | 16.3 (115.9) |
| No. of crystals used | | 1 | 1 |
| No. of Reflections Used: | Total | 5,736,096 | 6,279,665 |
| | Unique | 1,649,189 | 1,527,540 |
| Redundancy (outer shell) | | 3.48 (3.52) | 14.11 (3.54) |
| ***Refinement*** | | | |
| Resolution range of the diffraction data included in the refinement, Å | | 162-2.65 | 254-2.70 |
| $R_{work}/R_{free}$, % | | 20.2/25.2 | 21.8/27.9 |
| *No. of Non-Hydrogen Atoms* | | | |
| RNA | | 200,309 | 200,225 |
| Protein | | 90,976 | 90,976 |
| Ions (Mg, K, Zn, Fe) | | 2,827 | 2,863 |
| Waters | | 4,369 | 5,052 |
| *Ramachandran Plot* | | | |
| Favored regions, % | | 91.86 | 91.80 |
| Allowed regions, % | | 7.85 | 7.98 |
| Outliers, % | | 0.29 | 0.22 |
| *Deviations from ideal values (RMSD)* | | | |
| Bond, Å | | 0.008 | 0.009 |
| Angle, degrees | | 1.380 | 1.466 |
| Chirality | | 0.057 | 0.060 |
| Planarity | | 0.007 | 0.008 |
| Dihedral, degrees | | 17.342 | 17.871 |
| Average B-factor (overall), Å$^2$ | | 59.2 | 53.3 |

#### II. SUPPLEMENTARY FIGURES

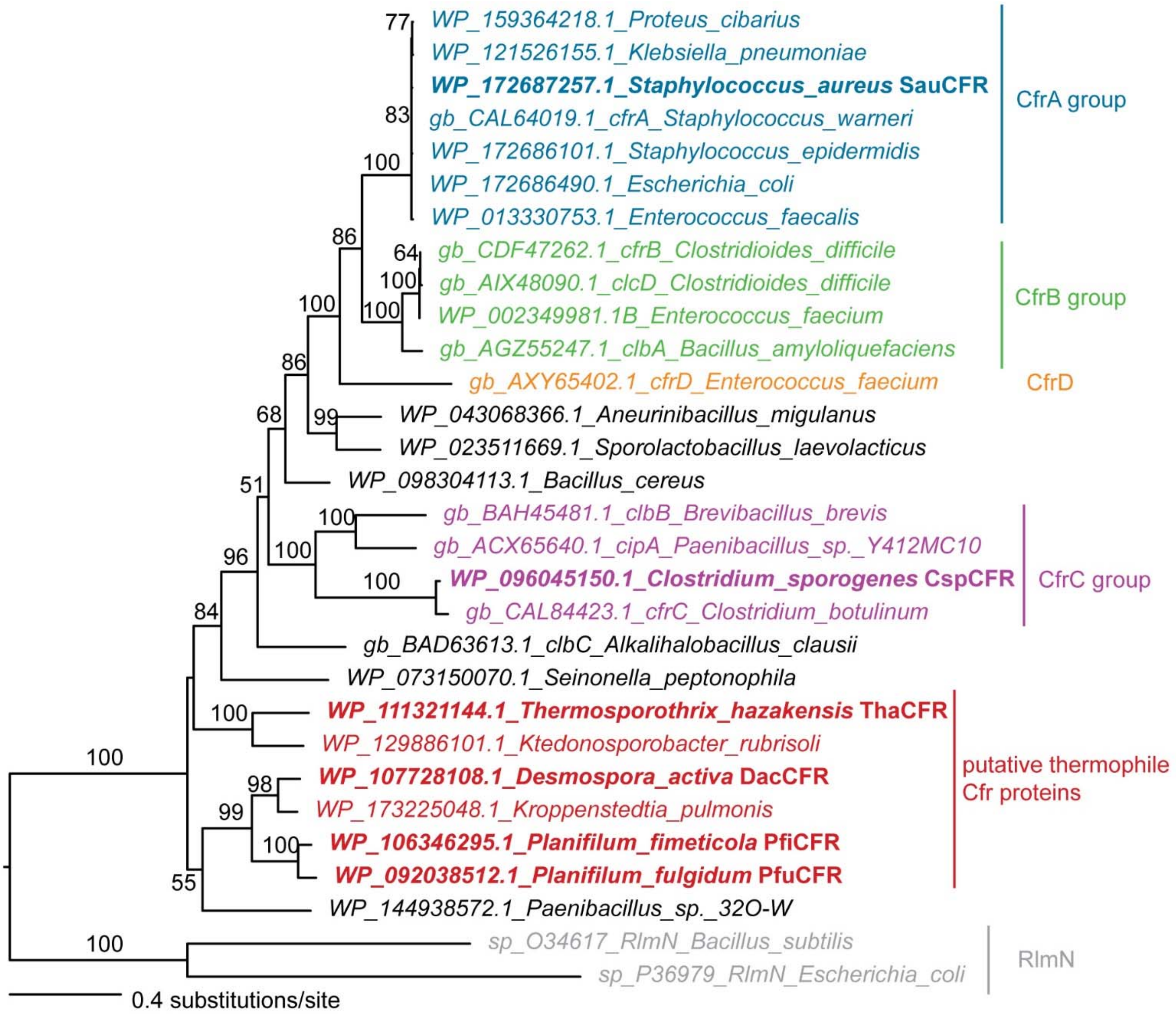


**Supplementary Figure 1 | Maximum likelihood phylogeny of Cfr-type proteins.** The well-studied reference Cfr-methyltransferase from *Staphylococcus aureus* is highlighted in blue. The cloned and tested Cfr-like proteins from moderately thermophilic bacteria and *S. aureus* are highlighted in bold. Branch lengths are proportional to the number of substitutions according to the lower-left key. Numbers on branches are IQTree ultrafast bootstrap support percentages. Unlabelled branches are those with bootstrap support values below 50%.

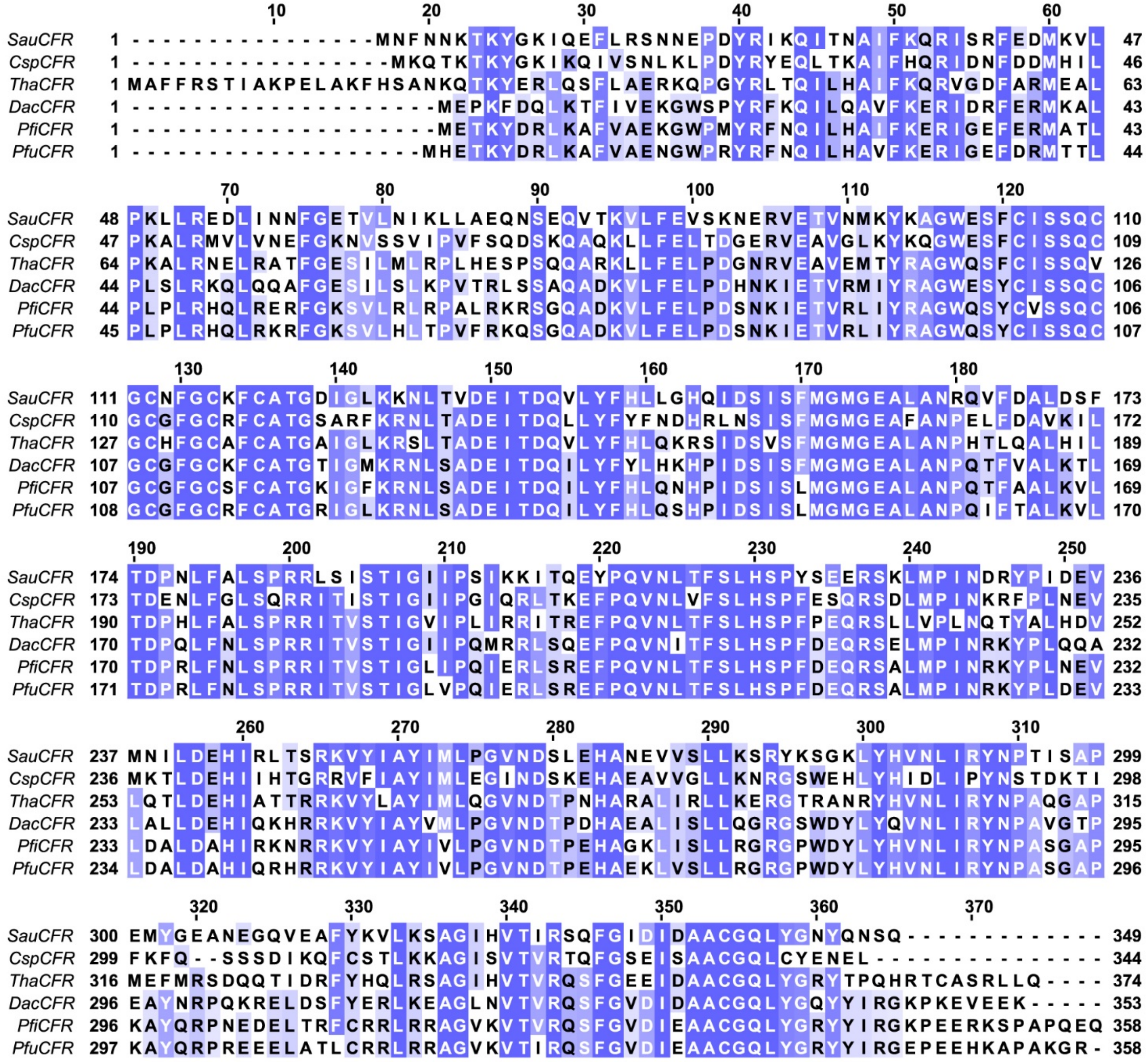


**Supplementary Figure 2 | Multiple sequence alignment of Cfr-like A2503-C8-methyltransferases from several bacterial species.** Multiple sequence alignment of well-studied Cfr-methyltransferase from mesophilic bacterium *Staphylococcus aureus* (SauCFR) with the putative Cfr-like proteins from moderately thermophilic *Clostridium sporogenes* (CspCFR)[1], *Thermosporothrix hazakensis* (ThaCFR)[2], *Desmospora activa* (DacCFR)[3], *Planifilum fimeticola* (PfiCFR)[3,4], and Planifilum fulgidum (PfuCFR)[3,4].

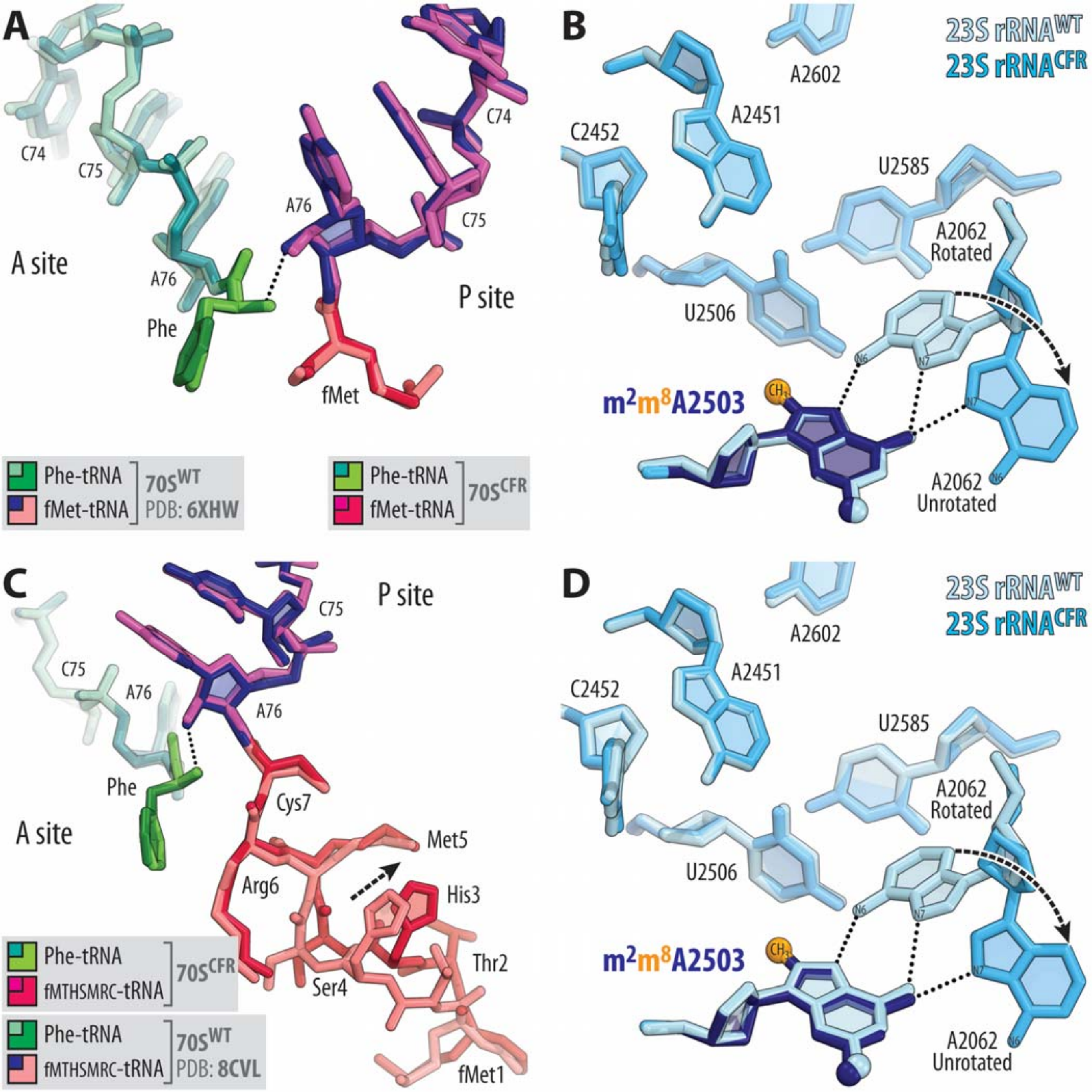


**Supplementary Figure 3 | Comparison of the structures of Cfr-modified and wild-type 70S ribosomes.** (**A, C**) Superpositioning of the previously reported structures of WT 70S ribosome containing aminoacylated Phe-tRNA$^{Phe}$ in the A site and either fMet-tRNA$_i^{Met}$ (A, PDB entry 6XHW[5]) or fMTHSMRC-peptidyl tRNA$_i^{Met}$ (C, PDB entry 8CVL[6]) in the P site with the new structures of precisely the same complexes containing Cfr-methylated nucleotide A2503 of the 23S rRNA. All structures were aligned based on domain V of the 23S rRNA. (**B, D**) Comparisons of the positions of key 23S rRNA nucleotides around the PTC in the same structures. Nucleotides of the Cfr-modified and unmodified ribosomes are shown in blue and light blue, respectively. The Cfr-modified A2503 residue is highlighted in navy, with the C8-methyl group shown in orange. *E. coli* nucleotide numbering is used. H-bonds are shown with dotted lines.

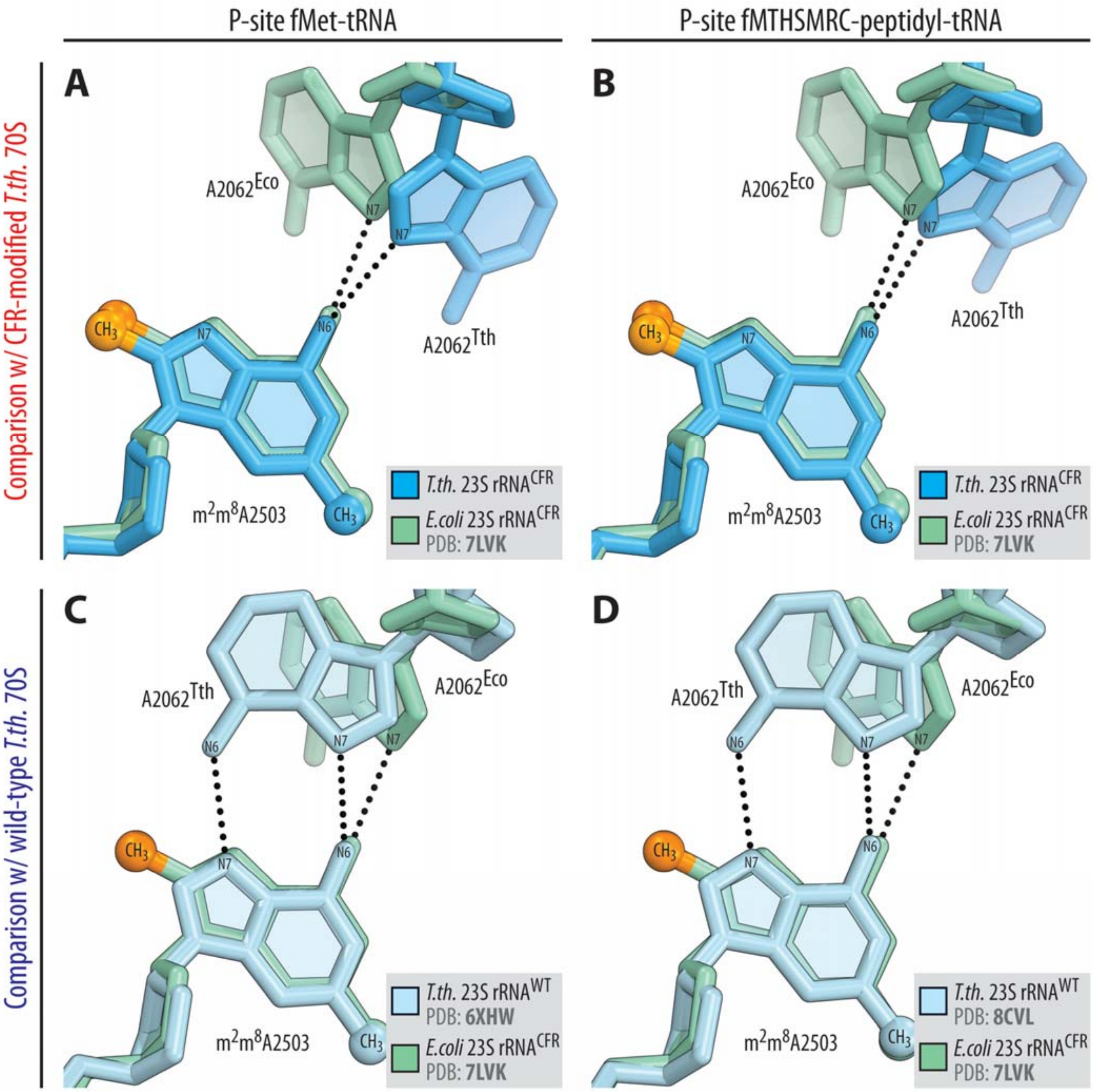


**Supplementary Figure 4 | Comparison of the structures of Cfr-modified ribosomes from *E. coli* and *T. thermophilus*.** (**A-D**) Superpositioning of the previous structure of Cfr-methylated *Escherichia coli* 50S ribosomal subunit (light teal, all panels, PDB entry 7LVK[7]) with the new structures of Cfr-methylated (A, B, blue) or wild-type (C, D, light blue) *Thermus thermophilus* 70S ribosome carrying Phe-tRNA$^{Phe}$ in the A site and either fMet-tRNA$_i^{Met}$ (A, C) or fMTHSMRC-peptidyl- tRNA$_i^{Met}$ (B, D) in the P site. Note that the C8-methylation does not affect the overall position of nucleotide A2503 in the Cfr-modified ribosome.

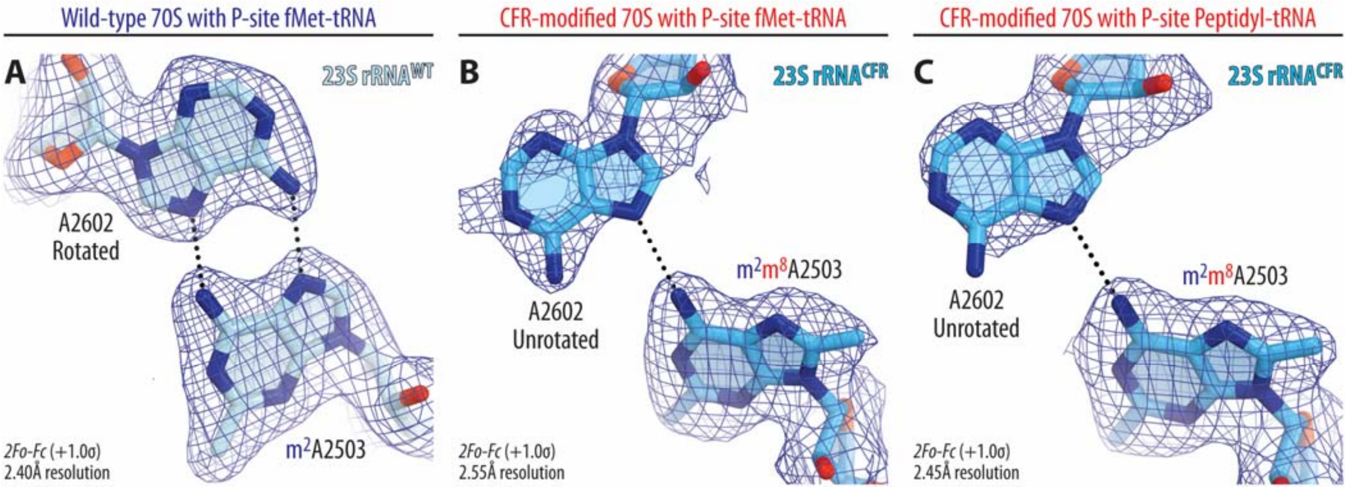


**Supplementary Figure 5 | Electron density maps of 23S rRNA in wild-type and Cfr-modified *T. thermophilus* 70S ribosome.** $2F_o\text{-}F_c$ electron difference Fourier maps (blue mesh) of A2062 and A2503 residues of 23S rRNA in the wild-type (**A**) or Cfr-modified (**B, C**) *T. thermophilus* 70S ribosome carrying aminoacylated Phe-$tRNA^{Phe}$ in the A site and either fMet-$tRNA_i^{Met}$ (A, B) or fMTHSMRC-peptidyl- $tRNA_i^{Met}$ (C) in the P site. The structure and the electron density map of the wild-type ribosome complex (A) are from PDB entry 6XHW[5]. Carbon atoms are colored light blue for the C8-unmethylated A2503 (A) and blue for the Cfr-modified A2503 (B, C); nitrogens are dark blue; oxygens are red.

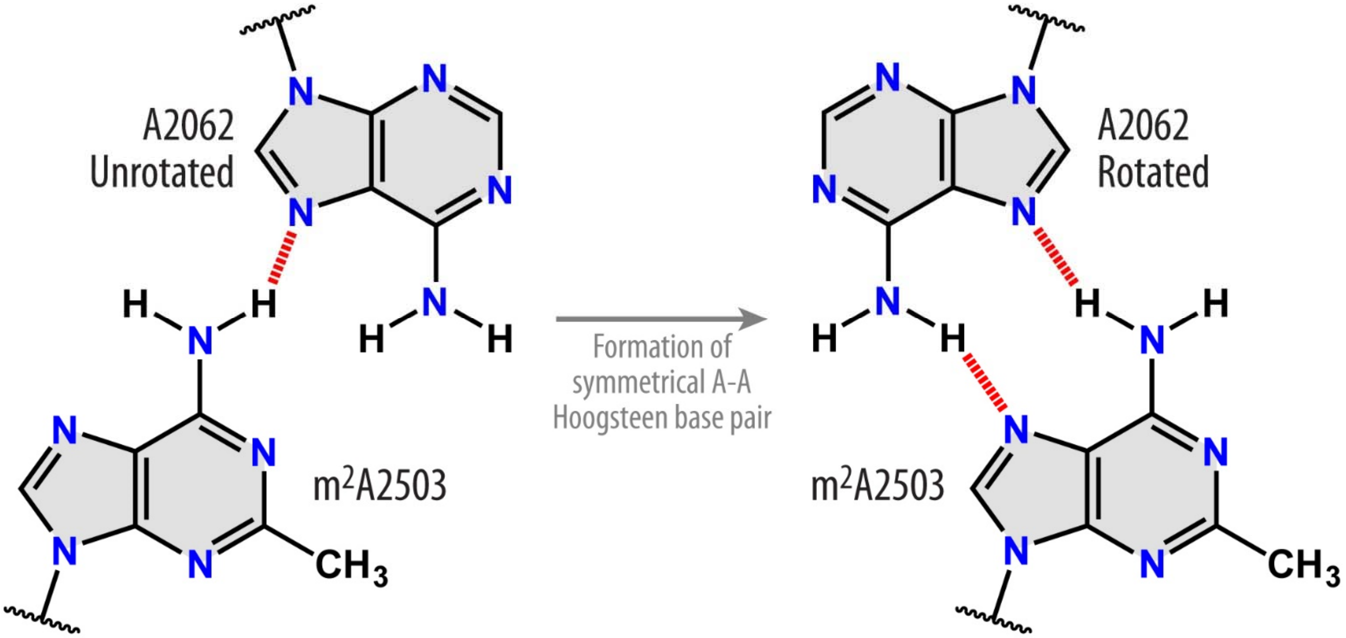


**Supplementary Figure 6 | Schematic diagram of H-bond rearrangement between nucleotides A2062 and A2503 of the 23S rRNA upon Hoogsteen base pair formation.** Note that the formation of a symmetric trans A-A Hoogsteen A2503-A2062 base pair requires the N7-atoms of both adenines to be deprotonated in order to serve as H-bond acceptors of the N6-protons of the base-paired nucleotide.

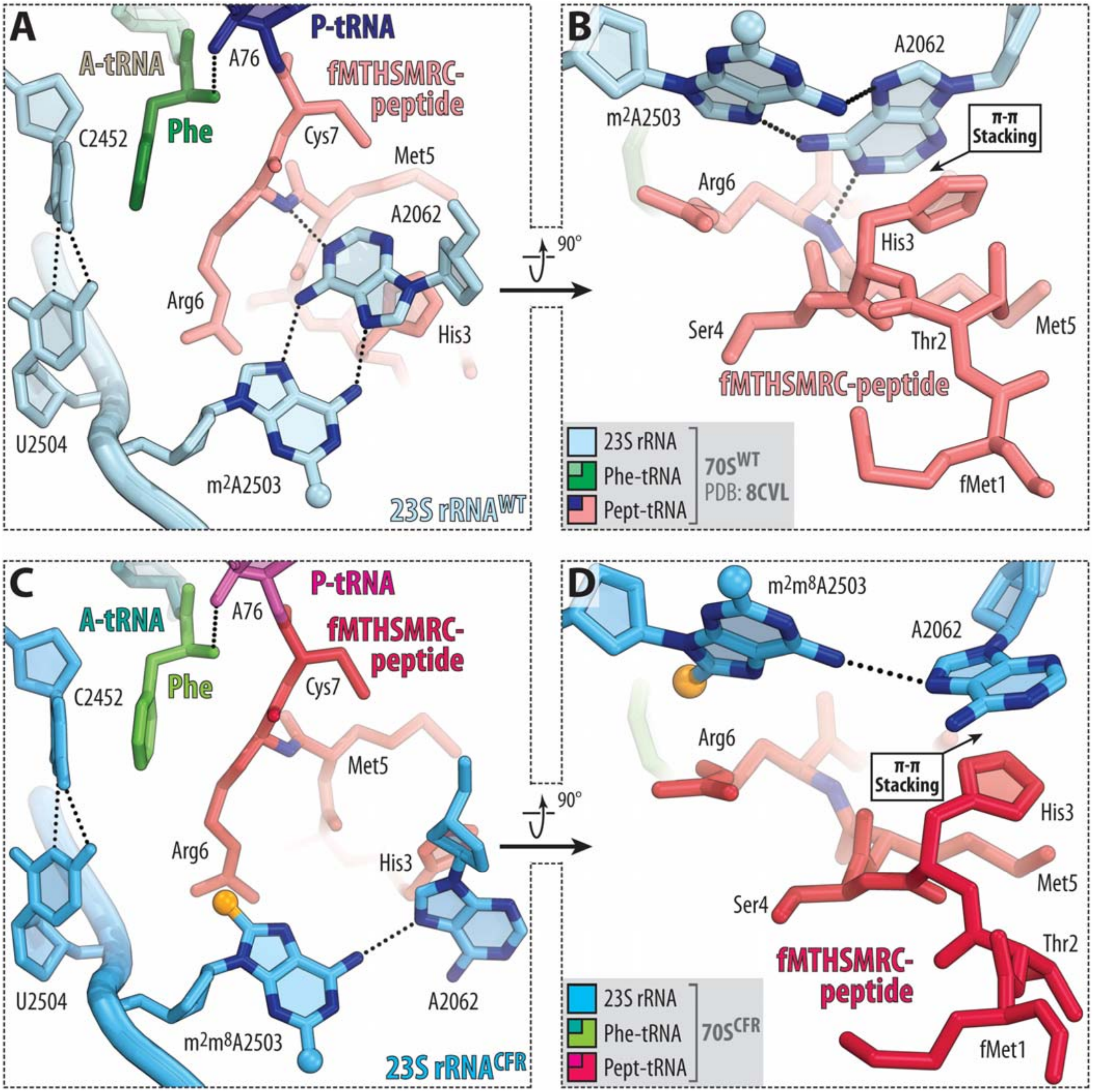


**Supplementary Figure 7 | Interactions of fMTHSMRC-peptidyl-tRNAs with wild-type and Cfr-modified *T. thermophilus* 70S ribosome.** Close-up views of the aminoacyl and peptidyl moieties of A-site Phe-tRNA$^{Phe}$ and P-site fMTHSMRC-tRNA$_i^{Met}$ in the wild-type (**A, B**, PDB entry 8CVL[6]) or Cfr-modified (**C, D**) *T. thermophilus* 70S ribosome. H-bonds are shown by black dotted lines. Stacking interactions between the aromatic side chain of His3 of fMTHSMRC-peptidyl-tRNA and A2062 nucleobase of the 23S rRNA are shown by black arrow.

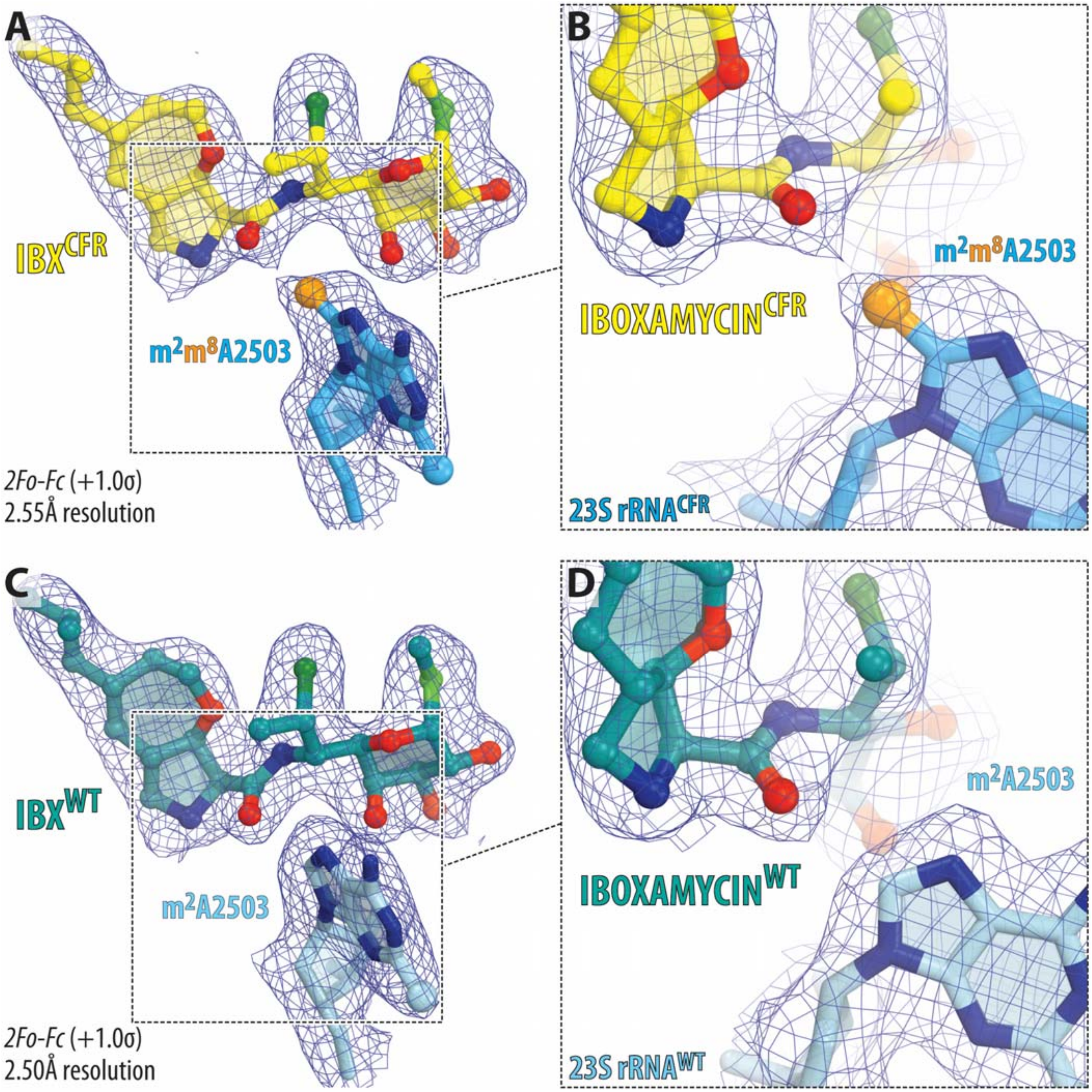


**Supplementary Figure 8 | Comparison of electron density maps of iboxamycin (IBX) in complex with Cfr-modified and wild-type *T. thermophilus* 70S ribosome.** (**A-D**) $2F_o$-$F_c$ electron density maps (blue mesh) contoured at 1.0σ of IBX in complex with Cfr-modified (A, B, yellow) or wild-type (C, D, teal) *T. thermophilus* 70S ribosome. The C8-methyl group of $m^2m^8$A2503 is highlighted in orange. The structure and the electron density map of IBX in complex with wild-type 70S ribosome (C, D) are from PDB entry 7RQ8[8]. Carbon atoms are colored light blue for the C8-unmethylated A2503 (A) and blue for the Cfr-modified A2503 (B, C); nitrogens are dark blue; oxygens are red.

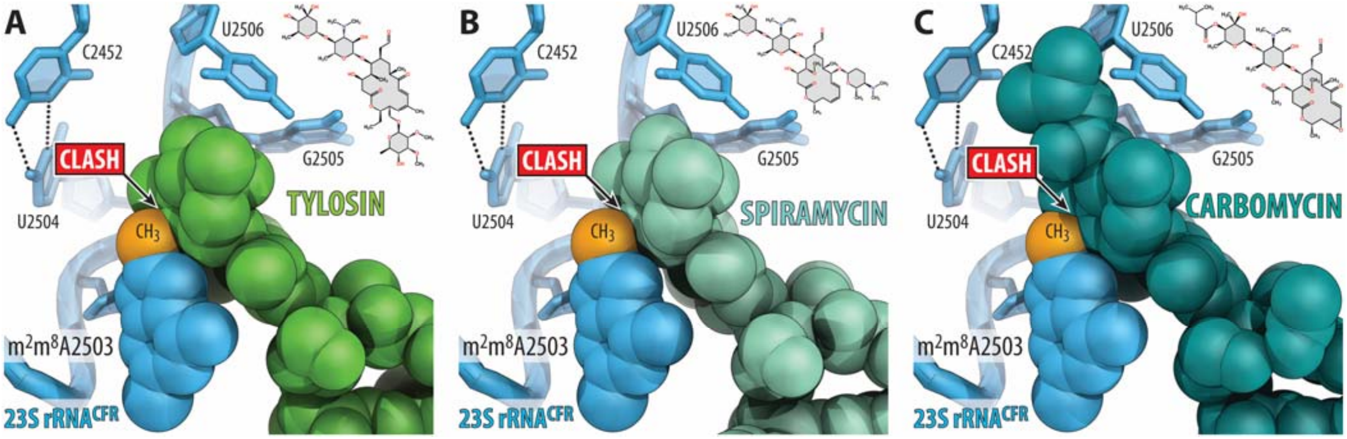


**Supplementary Figure 9 | Structural basis for the Cfr-mediated resistance to 16-membered macrolides.** (**A-C**) Superposition of the structures of Cfr-modified *T. thermophilus* 70S ribosome containing C8-methylated A2503 residue in the 23S rRNA (blue) with the previously reported structures of 16-membered macrolides, such as tylosin (A, green, PDB entry 1K9M[9]), spiramycin (B, light teal, PDB entry 1KD1[9]), or carbomycin (C, teal, PDB entry 1K8A[9]) in complex with *H. marismortui* 50S ribosomal subunit. Note that the highlighted in orange C8-methyl group of $m^2m^8$A2503 physically interferes with the binding of 16-membered macrolides.

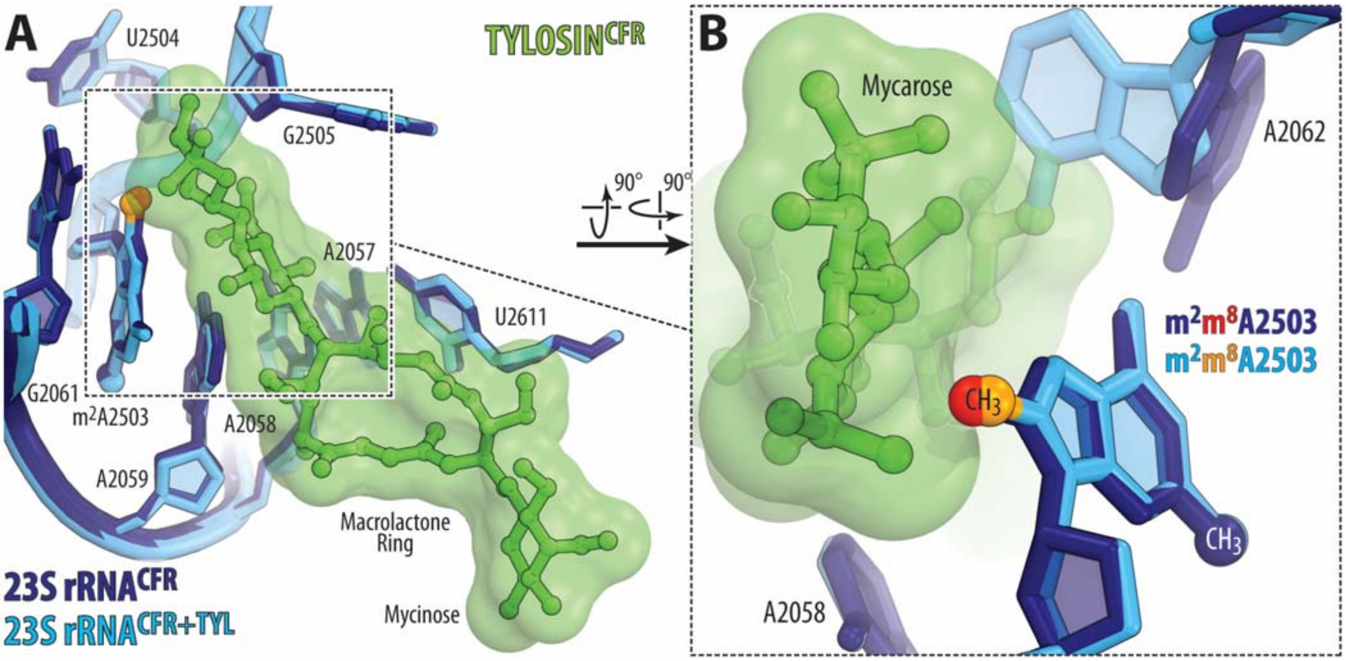


**Supplementary Figure 10 | Binding of tylosin to the Cfr-modified ribosome does not affect A2503 position.** (**A, B**) Superposition of the structures of Cfr-modified 70S ribosome containing an $m^2m^8$A2503 residue in the presence and absence of tylosin (green). Note that the position of $m^2m^8$A2503 residue is almost identical in the two structures.
